# In silico, in vitro, and in vivo models reveal EPHA2 as a target for decreasing inflammation and pathological endochondral ossification in osteoarthritis

**DOI:** 10.1101/2022.06.12.495737

**Authors:** Mauricio N. Ferrao Blanco, Raphaelle Lesage, Nicole Kops, Niamh Fahy, Fjodor T. Bekedam, Athina Chavli, Yvonne M. Bastiaansen-Jenniskens, Liesbet Geris, Mark G. Chambers, Andrew A. Pitsillides, Roberto Narcisi, Gerjo J.V.M. van Osch

## Abstract

Low-grade inflammation and pathological endochondral ossification are processes underlying the progression of osteoarthritis, the most prevalent joint disease worldwide. In this study, data mining on publicly available transcriptomic datasets revealed EPHA2, a receptor tyrosine kinase associated with cancer, to be associated with both inflammation and endochondral ossification in osteoarthritis. A computational model of cellular signaling networks in chondrocytes predicted that in silico activation of EPHA2 in healthy chondrocytes increases inflammatory mediators and triggers hypertrophic differentiation, the phenotypic switch characteristic of endochondral ossification. We then evaluated the effect of inhibition of EPHA2 in cultured human chondrocytes isolated from individuals with osteoarthritis and demonstrated that inhibition of EPHA2 indeed reduced inflammation and hypertrophy. Additionally, systemic subcutaneous administration of the EPHA2 inhibitor ALW-II-41-27 attenuated joint degeneration in a mouse osteoarthritic model, reducing local inflammation and pathological endochondral ossification. Collectively, we demonstrate that pharmacological inhibition of EPHA2 with ALW-II-41-27 is a promising disease-modifying treatment that paves the way for a novel drug discovery pipeline for osteoarthritis.

## Introduction

Osteoarthritis (OA) stands as the most widespread and debilitating musculoskeletal condition worldwide, characterized by progressive joint alterations leading to pain and functional limitations [1, 2]. In OA, joint inflammation ensues, resulting in the loss of cartilage lining the joint surface and the formation of bony outgrowths known as osteophytes at the joint edges. Current pharmacological treatments mainly focus on symptom management, primarily using pain relievers that do not impede disease advancement. Therefore, there is an imperative need to discover drugs capable of modifying OA progression.

Chondrocyte hypertrophy, a phenotype preceding cartilage calcification and eventual replacement by bone, plays a crucial role in OA pathogenesis. Under normal circumstances, articular chondrocytes resist hypertrophic changes. However, in OA, the onset of hypertrophic differentiation accelerates cartilage breakdown [3]. Similarly, joint margins are established through the growth of an initial cartilage template that undergoes endochondral ossification and is replaced by bone [4]. The inflammatory environment in OA, characterized by synovitis, involves heightened macrophage activation and the release of inflammatory cytokines like tumor necrosis factor-alpha (TNF-α) [5, 6]. Activation of inflammatory signaling pathways not only leads to cartilage matrix degradation but also triggers chondrocyte hypertrophy [7, 8].

Hence, we propose that targeting a key regulator governing both chondrocyte hypertrophy and inflammation could serve as an effective therapeutic approach for OA.

## Methods

### *In silico* simulations

A computational model of the intracellular signaling pathways regulating articular chondrocyte phenotypes was leveraged and completed with information about EPHA2 [9, 10]. In silico experiments involved setting targeted variables to specific values (0 for inhibition, 1 for activation) for 1,000 computing steps, followed by allowing variables to evolve freely until a new stable state was reached, simulating a bolus treatment effect. This process was repeated 100 times, and outcomes were averaged to compute final profiles and standard deviations. Four perturbations were applied: (1) EPHA2 activation, (2) pro-inflammatory cytokine activation, (3) combined activation of EPHA2 and pro-inflammatory cytokines, and (4) EPHA2 blockade with pro-inflammatory cytokine activation. The model and associated code are available via the following GitHub repository: https://github.com/Rapha-L/Virtual_Chondrocyte_for_EPHA2_study.

### Evaluation of ALW-II-41-27 in OA chondrocytes

*In vitro* validation was conducted using chondrocytes isolated from human articular cartilage obtained from OA donors (2 females, 1 male, aged 61, 64, and 69 years). Cartilage was obtained with implicit consent as waste material from patients undergoing total knee replacement surgery, approved by the medical ethical committee of the Erasmus MC, University Medical Center, Rotterdam (protocol number MEC-2004-322). Cartilage chips were subjected to protease and collagenase B digestion to isolate chondrocytes, which were then expanded in monolayer culture. For redifferentiation, chondrocytes were cultured in a 3D alginate bead model [11, 12]. After two weeks, cells were treated with 10 μM of ALW-II-41-27, vehicle (DMSO), and/or TNF-α for 24 hours. Medium and alginate beads were harvested for further analyses.

### Animal model

All animal experimentation procedures were conducted in compliance with the Animal Ethical Committee of Erasmus University Medical Center (License number AVD101002015114, protocol number 16-691-06). Twelve-week-old male C57BL/6 mice (C57BL/6J0laHsd, 27.01 g ± 2.05 g; Envigo, Cambridgeshire, UK) were group-housed in individually ventilated cages and maintained on a 12-hour light/dark cycle with unrestricted access to standard diet and water at the Experimental Animal Facility of the Erasmus MC. Mice were randomly assigned to two experimental groups (N=8 per group): Control and ALW-II-41-27-treated mice.

For all procedures, mice were anesthetized using 3% isoflurane/0.8 L O2/min (Pharmachemie BV, Haarlem, the Netherlands). Osteoarthritis (OA) was induced unilaterally by intra-articular injections of 60 μg Monoiodoacetate (MIA) (Sigma-Aldrich, St. Louis, USA) in 6 μl of saline (0.9% NaCl; Sigma-Aldrich) on day 0. Injections were administered following a 3-4 mm dermal incision made to the right knee at the height of the patellar tendon, using a 50 μl syringe (Hamilton, Bonaduz, Switzerland) and 30G needle (BD Medical, New Jersey, USA).

ALW-II-41-27 was administered via Alzet micro-osmotic pumps (Durect Corporation, CA, USA) model 1004, with delivery rates of 0.11 μl/hour, implanted subcutaneously on the back of the mice, slightly posterior to the scapulae, immediately after the intra-articular injections. Osmotic pumps were filled with dimethyl sulfoxide: polyethylene glycol alone (55:45 ratio, vehicle-treated group, N=8 mice) or containing 6 mg of ALW-II-41-27 dissolved in vehicle (treated group, N=8 mice), resulting in a dose of 6.6 μg/hour. A third group of N=8 mice received osmotic pumps delivering a dose of 1.7 μg/hour of ALW-II-41-27. A third group of N=8 mice received the implantation of osmotic pumps delivering a dose of 1.7 μg/ hour of ALW-II-41-27. In the figures 5 and 6 we report the 6.6 μg/ hour dose of ALW-II-41-27. Synovial thickness, Krenn score, and osteophyte size for all groups can be found in supplementary figure 6. Mice were euthanized in accordance with Directive 2010/63/EU by cervical dislocation under isoflurane anesthesia 14 days following MIA injection. Knee specimens were fixed in 4% formalin (v/v) for 1 week, decalcified in 10% EDTA for 2 weeks, and embedded in paraffin. Coronal sections of 6 μm were cut for analysis.

### Data and statistical analyses

Statistical analysis was conducted using GraphPad Prism 9.0 and IBM SPSS 24 (IBM). Each *in vitro* experiment comprised a minimum of 3 biological replicates and was replicated using cells obtained from 3 donors. The *in vivo* study was designed to ensure equal-sized groups, employing randomization and blinded analyses. The stated group size represents the number of independent values used for statistical analysis. Sample size for the *in vivo* study was determined based on the distribution of weight over the hind limbs as read-out parameter. Based on previous studies, we consider an increase of 13% (standard deviation of 10%) in weight distribution on the affected limb in time in the therapy groups as relevant in our study [13]. Sample size calculation was performed with a statistical power of 80% and a significance level of 0.05, resulting in N=8.

For statistical analysis, a linear mixed model with Bonferroni’s multiple comparisons test was utilized.

## Results

### *EPHA2* is as an inflammation-related gene upregulated in hypertrophic chondrocytes and osteoarthritic cartilage

To identify a new therapeutic target for OA linked with chondrocyte hypertrophy and inflammation, we conducted data analysis on two publicly available murine microarray datasets [14, 15]. Differentially expressed genes (DEGs) in the articular cartilage of mice with OA induced by destabilization of the medial meniscus (DMM) were compared to those from mice undergoing sham surgery, identifying OA-related genes. By intersecting this set with DEGs in the hypertrophic zone versus the proliferative zone of the mouse growth plate, we identified 172 genes differentially expressed in OA associated with chondrocyte hypertrophy. Among these, nine genes were associated with the gene ontology inflammatory response [16] (Figure 1 A & B). Subsequently, we examined the expression of these nine genes in a human microarray dataset obtained from OA and healthy articular cartilage [17], finding that three genes were also upregulated in human OA cartilage (Figure 1 C). While *GJA1* and *PTGS2* have been previously studied in the context of OA [18], *EPHA2*, a tyrosine kinase receptor, has an undisclosed role in OA. *Epha2* was 30-fold upregulated in OA versus sham mouse articular cartilage (adjusted p-value= 2E-02) [15], and 3-fold upregulated in human OA versus healthy cartilage (adjusted p-value= 1E-08) [17]. Additionally, *Epha2* was 19-fold higher in the hypertrophic compared to the proliferative zone of the murine growth plate (adjusted p-value= 1E-07) [14].

**Figure 1.**
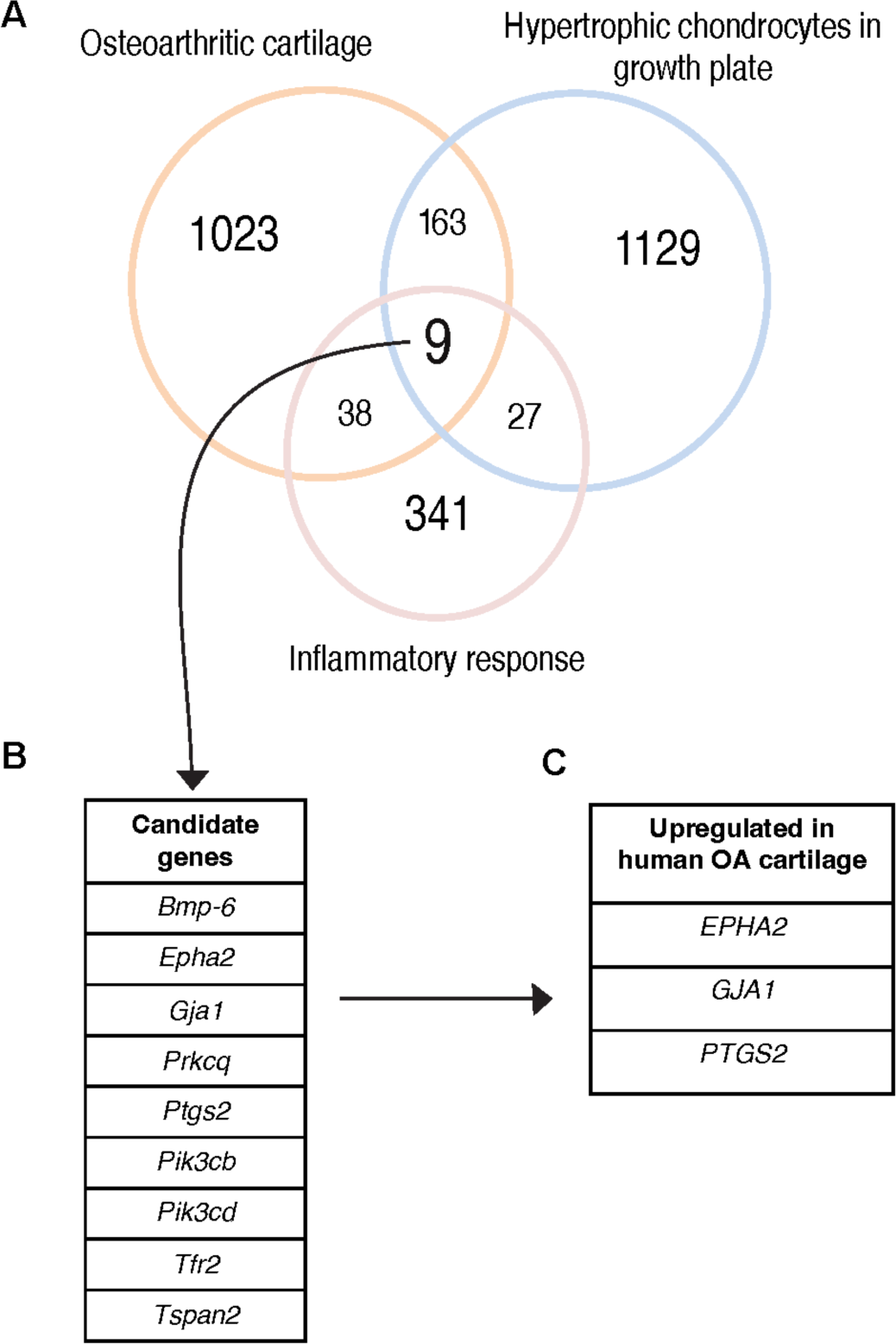
Identification of EPHA2 as a novel target for OA associated with inflammation and chondrocyte hypertrophy. (A) The Venn diagram illustrates the genes related to the inflammatory response, genes that were differentially regulated in murine osteoarthritic cartilage (DMM vs sham) and genes differentially expressed in the murine growth plate (proliferative vs hypertrophic zone). The number of genes in each dataset is represented, together with those that overlapped. (B) List of 9 targets that overlapped in the three databases. (C) Target genes upregulated in human OA compared with healthy articular cartilage. DEGs with a fold change greater than 3, and an adjusted p value lower than of 0.05, were considered for the analysis.

### EPHA2 triggers inflammatory signaling activation and hypertrophy in a virtual chondrocyte

To assess the role of EPHA2 on chondrocyte phenotype we utilized a computational model representing two cellular states: healthy and hypertrophic chondrocytes (Figure 2 A) [9, 19]. EPHA2 exhibited activity in the hypertrophic state but remained inactive in the healthy state (supplementary figure 1). Activation of EPHA2 led to a more pronounced transition of healthy chondrocytes towards the inflammatory hypertrophic state compared to activation of inflammatory cytokines (9% and 5% for ‘EPHA2+’ and ‘Inflammation+’, respectively; Figure 2 B & C). Consequently, both conditions prompted a decline in anabolic markers (collagen type II, Aggrecan: blue vs orange and yellow bars; Figure 2 D) and an increase in hypertrophic and inflammatory markers (Figure 2D & E). Full activation of both EPHA2 and inflammatory cytokines synergistically induced a complete transition of healthy chondrocytes towards the inflammatory hypertrophic state (’Inflammation+ EPHA2+’; Figure 2 C). Intriguingly, inhibiting EPHA2 (Inflammation+ EPHA2-) abolished this cellular state switching, accompanied by a significant reduction in the activity of hypertrophic and inflammatory entities (green vs violet and yellow bars; Figure 2 D & E). These computational findings substantiate the hypertrophic and inflammatory role of EPHA2 in chondrocytes.

**Figure 2.**
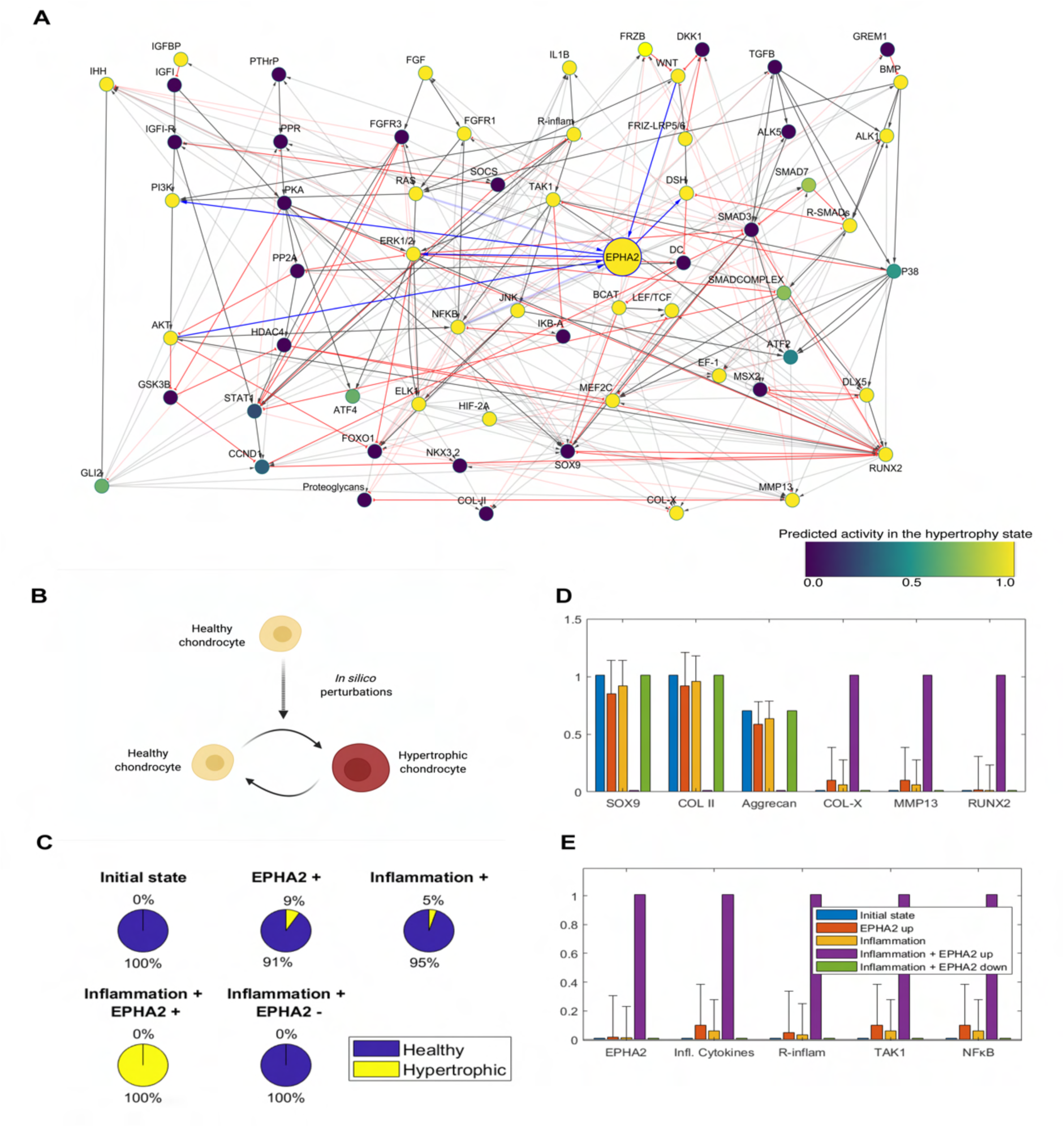
EPHA2 activation induces chondrocyte hypertrophy and inflammation in silico (A) Integration of EPHA2 in the regulatory network of articular chondrocytes. Solid lines denote protein interaction while modulation at the gene level is in dotted lines. Activating interactions are in grey with delta arrows and inhibitory interactions are in red with a half circle. Connections with EPHA2 are denoted in blue. All components of the model are represented with a variable that is named with capital letters. The variable represents neither the protein activation level nor the gene expression but a product of both (global activity). The color of the nodes in the network denotes the global activity of the variables in the hypertrophic state, with dark blue being 0 and yellow being 1. (B) Effect of gradually increasing the value of the EPHA2 input (from 0 to 1) in the scenario with EPHA2 activation and Inflammation, on the chondrocyte state transition. The initial state being the healthy (SOX9 +). (C) Activation of EPHA2 in the healthy state promotes the transition to a hypertrophic phenotype. Forced activation (+) or inhibition (-) of the entities from the initial healthy state is shown. Inflammation represents the activation of the variables related to inflammatory cytokines and their receptors, as input in the model. Pie charts represent the predicted percentage of perturbations leading to a transition to the hypertrophic phenotype or remaining in the healthy phenotype. (D) Average activity of the chondrogenic markers, being type II collagen, Aggrecan and SRY-box transcription factor (SOX9), and of hypertrophic markers, being Runt-related transcription factor 2 (RUNX2), Matrix Metallopeptidase (MMP13) and type 10 Collagen. (E) Average activity of EPHA2 and variables associated with inflammation. The bars represent the mean of the results of a hundred in silico experiments (+ standard deviations, +SD). There is no SD for the initial state (blue condition) as it denotes the starting point before the in silico perturbation is applied. Figure created with Cytoscape, MATLAB and Biorender.com.

### ALW-II-41-27 reduces human OA-derived chondrocyte inflammation

To investigate the potential of pharmacologically inhibiting EPHA2 to alleviate inflammation in OA, we utilized the tyrosine kinase inhibitor ALW-II-41-27, a type II kinase inhibitor known for its high selectivity for EPHA2 [20, 21]. Initially, we validated the ability of ALW-II-41-27 to reduce TNF-α-induced phosphorylation of EPHA2 in a dose-dependent manner (Figure 3 A). Notably, the concentration of 10 micromolars exhibited the highest efficacy in decreasing both EPHA2 phosphorylation and the expression of catabolic enzymes MMP1 and MMP13 in OA cartilage explants (supplementary figure 3). Subsequently, we examined the effects of ALW-II-41-27 at a concentration of 10 micromolars on TNF-α-treated human chondrocytes from OA donors (Figure 3 B). Treatment with ALW-II-41-27 significantly inhibited TNF-α-induced inflammatory responses, as evidenced by a reduction in the secretion of nitric oxide metabolites (Figure 3 C). Furthermore, ALW-II-41-27 administration mitigated the TNF-α-induced expression of the inflammatory cytokine interleukin (IL)-6, known to be a downstream target of TNF-α (Figure 2 D & E) [22]. Moreover, ALW-II-41-27 treatment countered the TNF-α-induced upregulation of the cartilage matrix-degrading enzymes MMP1 and MMP13 (Figure 3 F & G). These findings demonstrate the anti-inflammatory potential of ALW-II-41-27 in TNF-α-stimulated OA chondrocytes.

**Figure 3.**
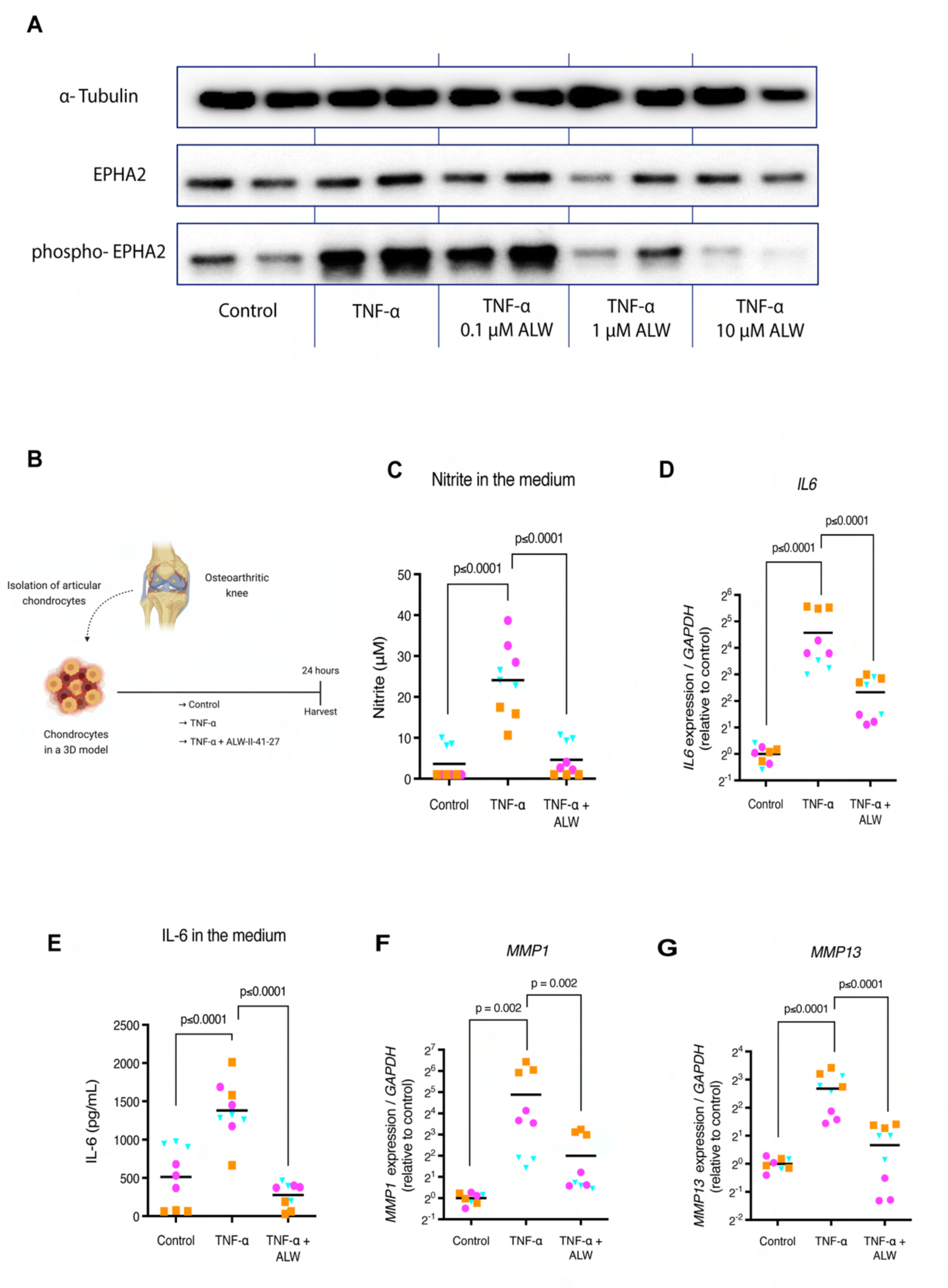
ALW-II-41-27 decreases TNF-α induced catabolism and inflammation in human chondrocytes. (A) ALW-II-41-27 decreases TNFα-induced phosphorylation of EPHA2 in a dose-dependent manner. (B) Experimental set-up to evaluate the anti-inflammatory capacity of ALW-II-41-27 (10 µM) in human OA chondrocytes cultured with 10 ng/mL TNF-α. (C) Evaluation of nitrite in the medium as a marker for inflammatory activity as determined by Griess reagent. (D, F-G) Gene expression of IL6, MMP1 and MMP13 determined by qPCR. The average of control, per donor, is set to 1. (E) IL-6 in the medium determined by ELISA. Experiments were performed in triplicate, with cells from three donors. Donors are represented with different colors and symbols: violet circles (donor 1), blue triangles (donor 2) and orange squares (donor 3). The horizontal line in the graphs represents the mean. Data were analyzed with the linear mixed model with Bonferroni’s multiple comparisons test. Figure created with Biorender.com.

### ALW-II-41-27 decreases chondrocyte hypertrophy

We then proceeded to evaluate whether pharmacological inhibition of EPHA2 with ALW-II-41-27 could impede hypertrophy in human OA-derived articular chondrocytes (Fig. 4 A). ALW-II-41-27 decreased *COL10A1* expression (Figure 4 B), suggesting a reduction in hypertrophy. Given that chondrocyte hypertrophy poses a significant obstacle to stable hyaline cartilage tissue engineering [23], we further investigated the effect of ALW-II-41-27 on cartilage tissue engineered constructs from mesenchymal stromal cells (MSCs), well-known to become hypertrophic and prone to ossify when implanted *in vivo* [24–27] (Figure 4 C). Gene expression analysis indicated that the addition of ALW-II-41-27 mitigated the hypertrophic markers *COL10A1* and *ALPL* (Figure 4 D & E). Likewise, when treated with ALW-II-41-27, the cells exhibited reduced deposition of type X Collagen (Figure 4 F and supplementary figure 4). Intriguingly, neither glycosaminoglycan nor type II Collagen deposition showed reduction, suggesting that ALW-II-41-27 primarily targeted hypertrophy.

**Figure 4.**
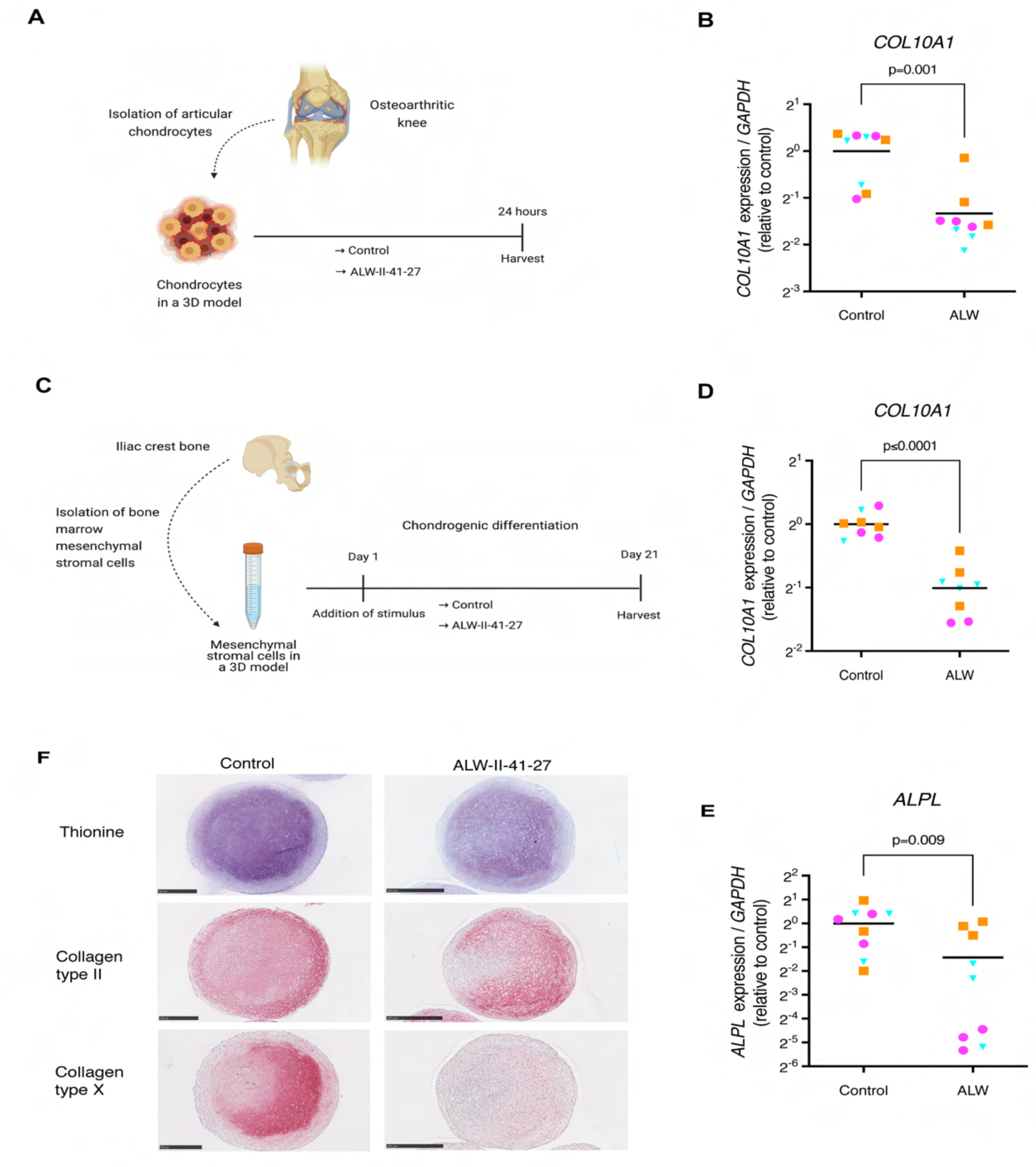
ALW-II-41-27 decreases chondrocyte hypertrophy. (A) Experimental set-up to evaluate the capacity of ALW-II-41-27 (10 µM) to modulate hypertrophy in human OA chondrocytes. (B) Gene expression of the hypertrophic marker COL10A1 in cultured OA chondrocytes determined by qPCR. (C) Experimental set-up to evaluate how ALW-II-41-27 (100 nM) affects hypertrophy in tissue engineered cartilage from MSCs. (D, E) Gene expression of COL10A1 and ALPL in MSC-generated cartilage determined by qPCR. (F) Histological analysis of tissue engineered cartilage derived from MSCs; Thionine staining showing glycosaminoglycans (violet), and immunohistochemistry of type II and type X collagen (red/pink). Experiments were performed with 3 replicates, for each of the three donors. Donors are represented with different colors and symbols: violet circles (donor 1), blue triangles (donor 2) and orange squares (donor 3). For gene expression analysis, the average of control replicates is set to 1 per donor. The horizontal line in the graphs represents the mean. For statistical analysis, the linear mixed model with Bonferroni’s multiple comparisons test was performed. Figure created with Biorender.com.

### ALW-II-41-27 treatment reduces joint inflammation and pathological endochondral ossification *in vivo*

EPHA2 inhibition reduced hypertrophy and inflammation *in silico* and its pharmacological inhibition with ALW-II-41-27 in cell culture *in vitro* confirmed these results. Subsequently, we sought to assess the potential therapeutic efficacy of ALW-II-41-27 in vivo using a mouse model of joint pain and degeneration induced by intra-articular injection of monoiodoacetate (MIA) [28]. ALW-II-41-27 was administered subcutaneously with a controlled delivery system (Figure 5 A). Notably, the total body weight remained unaffected by ALW-II-41-27 administration throughout the 14-day study period, indicating no significant adverse effects on the general health of the animals (supplementary figure 5).

**Figure 5.**
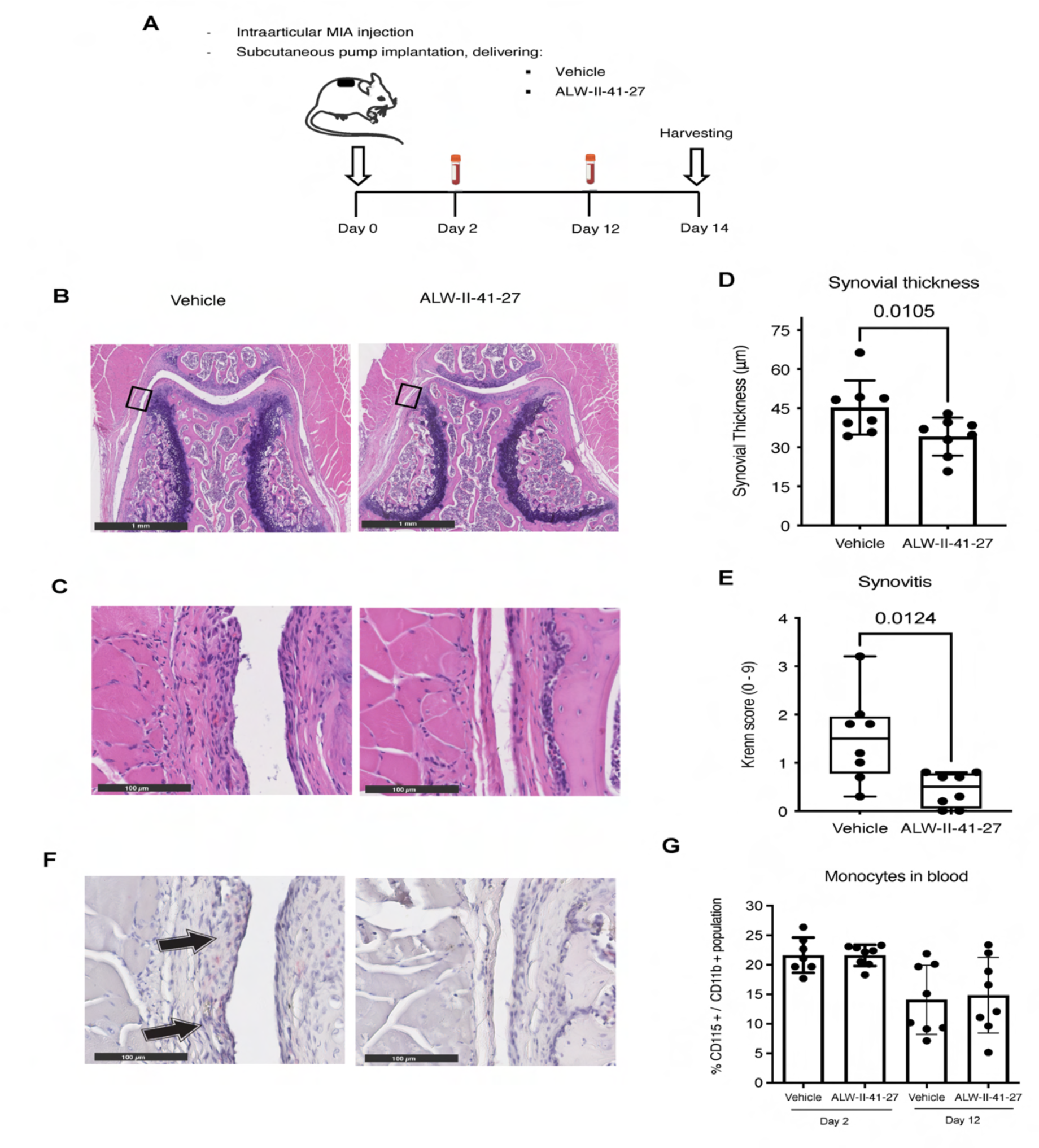
ALW-II-41-27 treatment attenuates joint inflammation. (A) Experimental set-up of the in vivo experiment. Intra-articular injection of monoiodoacetate (MIA, 60 µg in 6 μl of saline) was applied to the right knee of mice for each experimental group (N=8) to induce OA. An osmotic pump was implanted on the back of the mice, slightly posterior to the scapulae, which continuously delivered vehicle or ALW-II-41-27 in a dose of 6.6 µg/ hour. At day 2 and 12 peripheral blood was harvested. At day 14 mice were euthanized to assess the effects of ALW-II-41-27 on the degenerated joint. (B) Hematoxylin and Eosin staining of knees (patellofemoral region) from vehicle-treated and ALW-II-41-27-treated mice. Black square indicates the region of magnification for the image below (C) showing the synovial lining where synovial thickness and Krenn score were determined. (D) Synovial thickness is represented by the mean ± SD. (E) Krenn score illustrated with box-and-whiskers plots, with line indicating the median and error bars spanning maximum to minimum values. (F) Immunohistochemistry of F4/80 (pink) showing macrophages in the synovial lining. Arrows indicate positive staining. (G) Percentage of monocytes present in the peripheral blood of mice at day 2 and 12, respect to the myeloid cell population. In all graphs, each dot represents data of an individual mouse (N=8). For statistical analysis, the linear mixed model with Bonferroni’s multiple comparisons test was performed. Figure created with Biorender.com.

Treatment with ALW-II-41-27 led to a reduction in synovial membrane thickness and synovitis compared to vehicle-treated mice (Figure 5 B, C, D & E). Synovitis is governed by macrophages, the crucial regulators of OA progression and primary mediators of the inflammatory response [29, 30]. Macrophages were solely observed in the synovial lining of vehicle-treated mice, suggesting effective attenuation of joint inflammation (Figure 5 F). Moreover, there were no discernible alterations in peripheral blood monocyte levels at day 2 and 12 (Figure 5 G), indicating selective reduction of local joint inflammation without affecting systemic immune cells.

Despite joint pain being a significant OA symptom often associated with inflammation [31], no significant difference in weight distribution between limbs was detected between ALW-II-41-27-treated and vehicle-treated mice, suggesting no apparent effect on pain (supplementary Figure 6).

Cartilage degeneration, evidenced by proteoglycan loss, was observed in all mice irrespective of ALW-II-41-27 treatment (Figure 6 A & B), suggesting that ALW-II-41-27 was not able to rescue matrix degeneration *in vivo*. However, histological assessment revealed significantly smaller osteophytes, particularly at the lateral side of the patella, in ALW-II-41-27-treated mice (Figure 6 C & D). Additionally, type X collagen deposition was reduced in the knees of ALW-II-41-27-treated mice (Figure 6 E), suggesting effective targeting of endochondral ossification in the injured joint.

**Figure 6.**
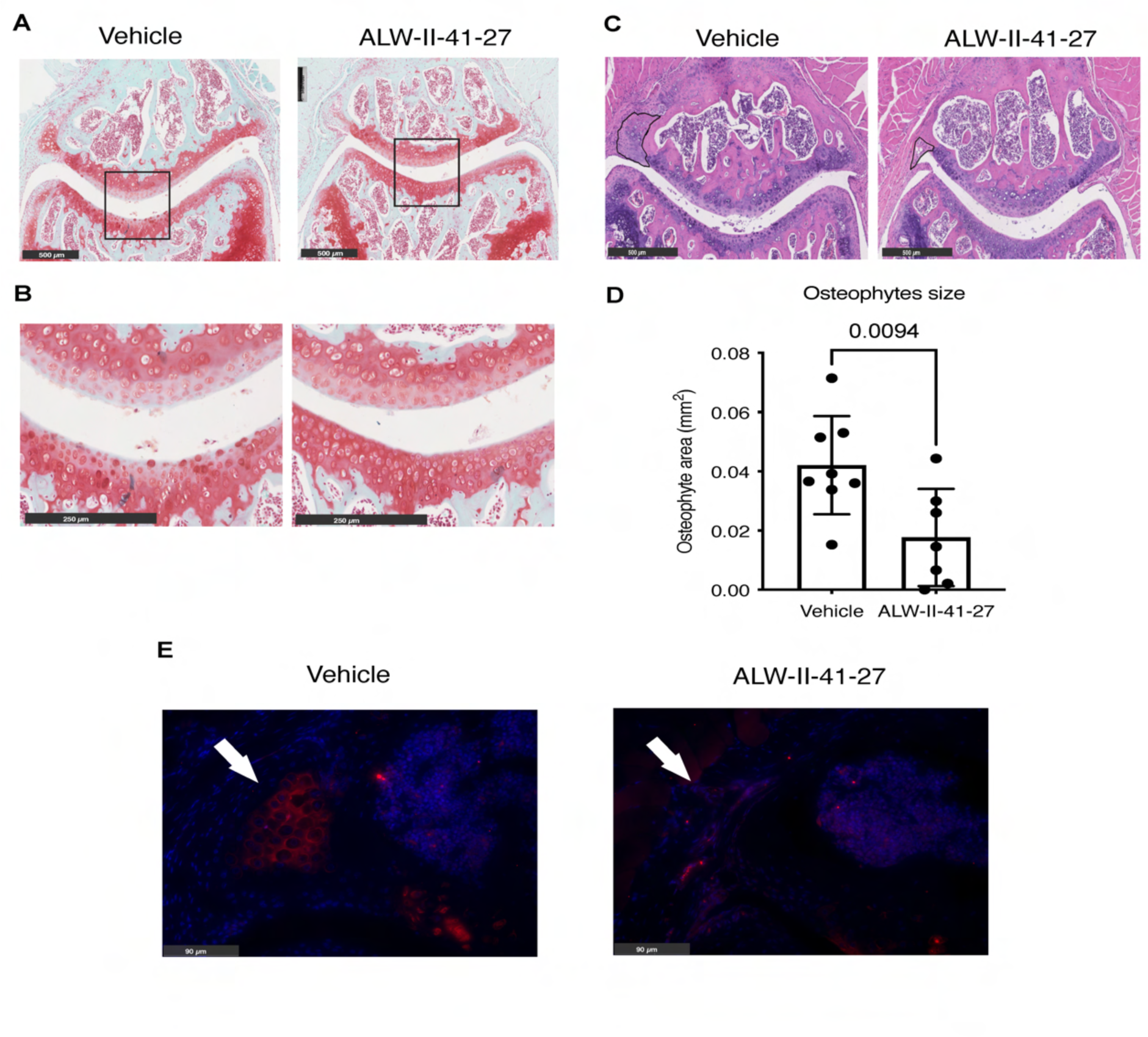
ALW-II-41-27 treatment attenuates pathological endochondral ossification (A) Safranin-O / Fast Green staining of vehicle-treated and ALW-II-41-27-treated mice knees with magnification of the patellofemoral region. Black square indicates the region of magnification for the image below (B) showing the central part of the patellofemoral articular cartilage. (C) Hematoxylin and Eosin stain of vehicle-treated and ALW-II-41-27-treated mice knees with magnification of the patellofemoral region. Osteophyte’s diameter is indicated with a black line in the lateral side of the patella. (D) Osteophyte area adjacent to the lateral side of the patella is represented by the mean ± SD. Each dot represents data of an individual mouse (n=8). For statistical analysis, the linear mixed model with Bonferroni’s multiple comparisons test was performed (E) Immunofluorescence of type X collagen (red) and DAPI (blue) in the lateral side of the patellofemoral region. Arrow indicates an osteophyte in the lateral side of the patella.

## Discussion

There is an urgent unmet need for effective therapies for OA patients. Here, we show that EPHA2 is a promising drug target for OA and we report the small molecule ALW-II-41-27 as a disease-modifying OA drug (DMOAD), specifically targeting inflammation and pathological endochondral ossification (Fig. 7A). To find targets associated with inflammation and chondrocyte hypertrophy we have used a particular sequence of studies that involved *in silico*, *in vitro* and *in vivo* analyses (Figure 7 B). For the i*n silico* analyses we leveraged previously published large gene expression dataset depositories and narrowed them down to one target of interest. The role of the identified target on inflammation and chondrocyte hypertrophy was further investigated through *in silico* experiments using a computational model of cellular signaling networks controlling chondrocyte phenotypes. These *in silico* studies served as the starting point for a series of *in vitro* experiments using different cell models, which were followed by an *in vivo* study. Our study illustrates the efficacy of this experimental approach in uncovering a novel target for specific biological processes in osteoarthritis While previous research has implicated EPHA2 as a key player in diseases like cancer [32–35] and irritable bowel disease [36], our study marks the first to underscore its significance in OA. Other tyrosine kinases, including fibroblast growth factor receptor (FGFR) 1, Fyn and vascular endothelial growth factor receptor (VEGFR), have been implicated to promote chondrocyte hypertrophy [37–40]. Our findings reveal that the tyrosine kinase EPHA2 not only contributes to hypertrophy but also to inflammation, making it a compelling target for mitigating pathological mechanisms in OA.

**Figure 7.**
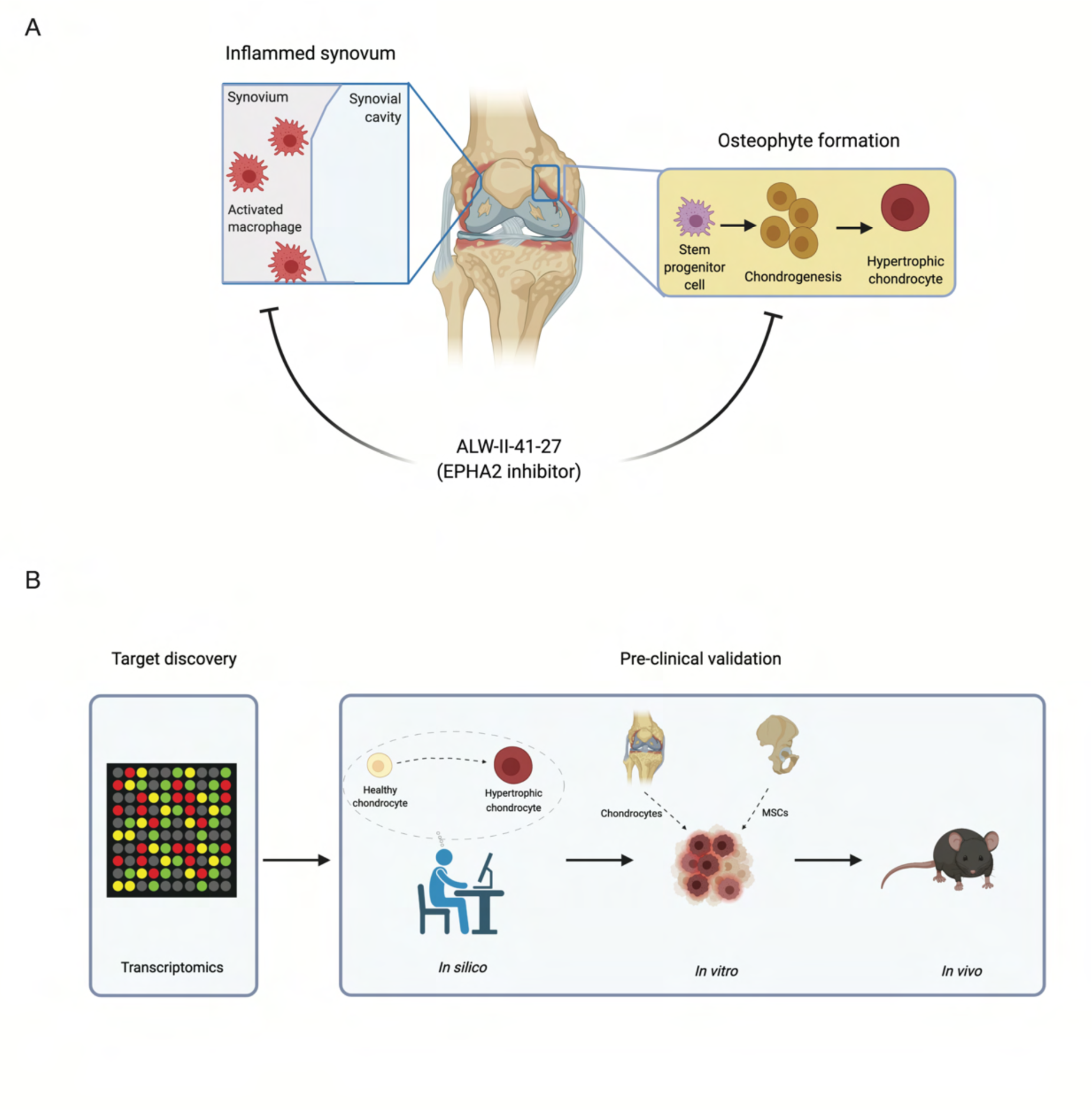
Graphical representation of main findings and experimental approach. (A) ALW-II-41-27 attenuates synovitis and osteophyte formation. (B) Drug discovery pipeline combining transcriptomic datasets, in silico, in vitro and in vivo models. Figure created with Biorender.com.

We have identified a promising compound, ALW-II-41-27, demonstrating potential as a disease-modifying osteoarthritic drug (DMOAD). To date, a wide variety of DMOADs targeting chondrocyte hypertrophy have been tested in experimental pre-clinical studies [41–43]. It is unclear, however, whether those agents have been subjected to further research. Regulatory agencies, such as the Food and Drug Administration (FDA) or the European Medicines Agency (EMA), have not yet approved any existing disease-modifying pharmacological intervention for OA [44]. Considering the pre-clinical data of ALW-II-41-27 to modulate OA pathogenesis and its extensive pharmacological analysis in other conditions [32–36], it is expected that clinical trials involving this compound may pose lower risks with a higher likelihood of success.

In addition to cartilage hypertrophy, inflammation of the synovium plays a crucial role in OA pathology [45]. Our results indicate that ALW-II-41-27 exhibits anti-inflammatory properties, aligning with its previously observed effects in a model of irritable bowel syndrome [36]. EPHA2 is not limited to chondrocytes but is also expressed in various other cell types found in the synovium, including fibroblasts, monocytes, and macrophages [46, 47]. Thus, the observed anti-inflammatory mechanism in our *in vivo* setting may also be linked to the action of ALW-II-41-27 on these cell types.

Our study has certain limitations. Treatment initiation coincided with OA induction in our research. This decision was influenced by the progressive nature of the disease, posing a challenge for a DMOAD to reverse extensive joint structural changes in end-stage OA. Hence, maximizing the pharmacological benefits of ALW-II-41-27 to reduce inflammation and osteophytosis might be more effective if administered during the earlier stages of the disease. Our study did not demonstrate that *in vivo* administration of ALW-II-41-27 prevented cartilage loss or pain in the MIA mouse model. Further investigation using alternative experimental animal models is warranted to determine whether ALW-II-41-27 specifically targets hypertrophy and inflammation, or if its effects extend to preventing cartilage degeneration and alleviating pain. Additionally, exploring the potential effects of ALW-II-41-27 on post-traumatic OA or other non-chemically induced forms of OA is essential.

In conclusion, our findings suggest that EPHA2 contributes to the pathogenesis of OA. The use of ALW-II-41-27 to inhibit EPHA2 showed promising results in mitigating inflammation and pathological endochondral ossification across various models, including *in silico* simulations, *in vitro* experiments with patient-derived cells, and a mouse model. These results underscore the potential of ALW-II-41-27 as a candidate drug for modifying the course of OA, warranting further investigation.

All authors approved the final version of the manuscript.

## Ethical statement

Animal experiments were approved by the medical ethical committee of the Erasmus MC, protocol EMC 16-691-06. Human articular cartilage was obtained with the approval of Erasmus MC, protocol MEC-2004-322 and mesenchymal stromal cells MEC-2014-16.

## Funding

This study was financially supported by the European Union’s Horizon 2020 research and innovation programme under the Marie Sklodowska-Curie grant agreement no. 721432 Carbon and the Reumafonds ReumaNederland grant number 18-1-202.

## Supplementary methods

Description of further procedures can be found in supplementary methods.

## Competing interest statement

The author(s) declared the following potential conflicts of interest with respect to the research, authorship, and/or publication of this article: M.G. Chambers is an employee of Eli Lilly but they did not fund this research.

## Supporting information

Supplementary Data_ListInteractions

ARRIVE guidelines

## Notes

https://github.com/Rapha-L/Virtual_Chondrocyte_for_EPHA2_study

### Competing Interest Statement

The authors have declared no competing interest.

### Summary of Updates

Update April 2024. Supplemental files and figures revised

