## Supplementary Figures for "In silico, in vitro, and in vivo models reveal EPHA2 as a target for decreasing inflammation and pathological endochondral ossification in osteoarthritis"

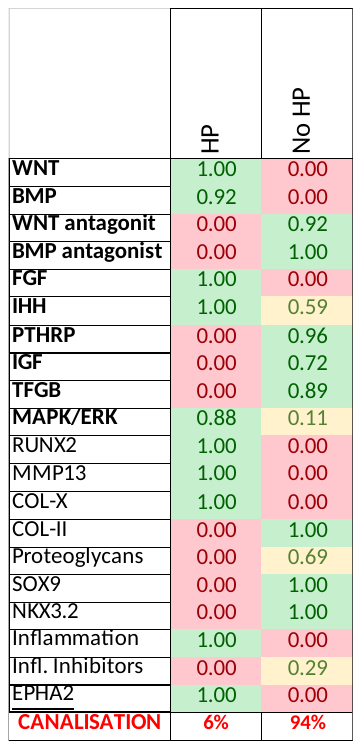

**Supplementary figure 1.** Summary of global predicted activity profiles for the two stable states emerging from the random initialization (Monte Carlo). The canalization indicates the chance of occurrence in the unconstrained system (i.e. the percentage of initial states, among 10.000, reaching the final state). Bold labels are used when the average activity of the pathway is displayed while regular labels are used for specifics single entities (i.e. variables). Values and colors reflect the global activity level, i.e. the product of the predicted gene expression and of the protein activation or repression level. HP= hypertrophic chondrocyte.

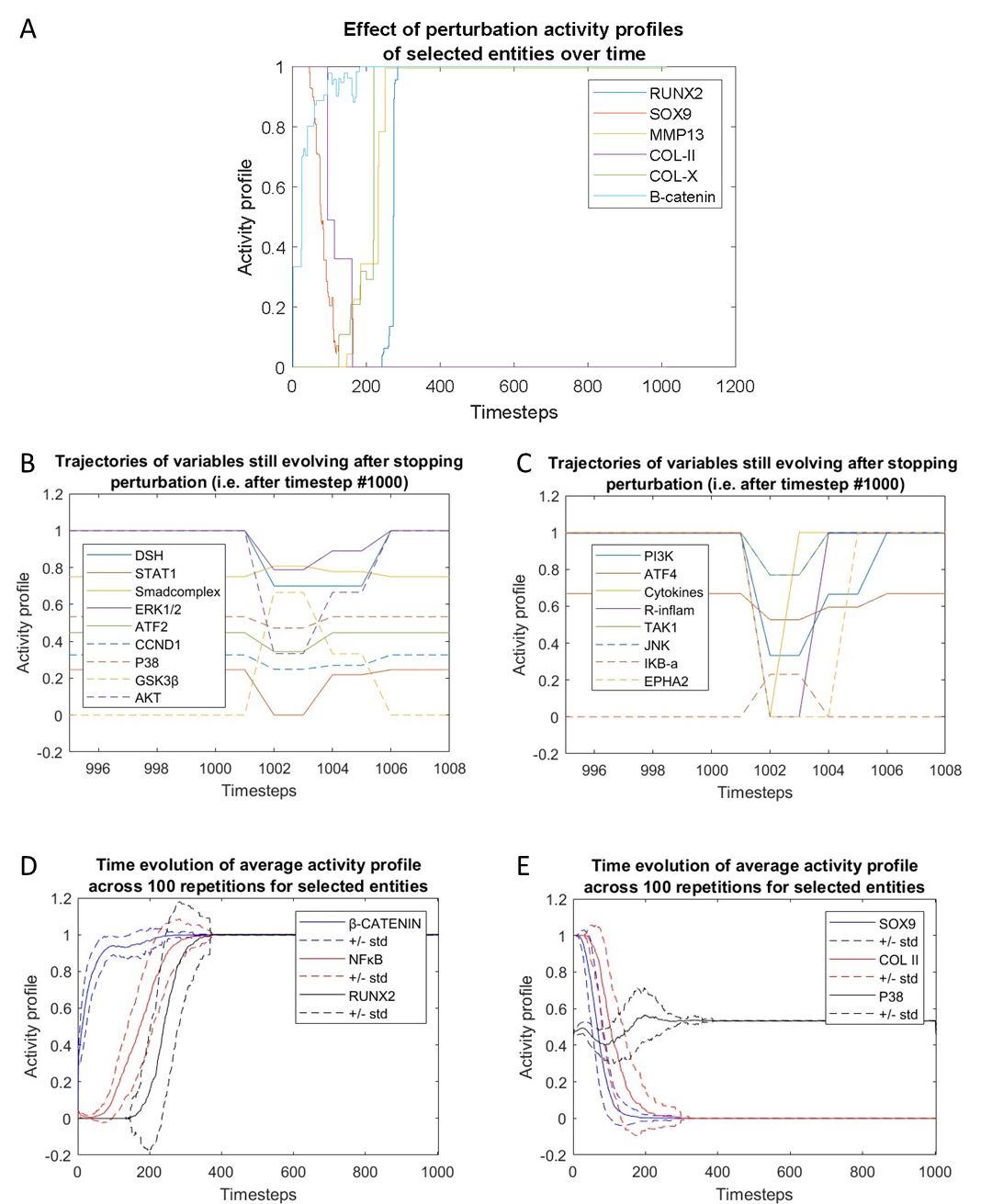

**Supplementary figure 2.** Evolution of variables over computational timesteps under perturbation. All figures reflect the effect of EPHA2 activation with input of inflammatory cytokines (both activities set to 1) on a healthy chondrocyte, in silico. Input perturbations are normally imposed for 1000 timesteps and then released until a stable state is reached. Perturbations are repeated 100 times since various outcomes may emerge due to stochasticity. Panel A-E depict example of trajectories for repetition n°66 of the perturbation. (A) Example of trajectories for RUNX2, SOX9, MMP13, COl2, COL10 and Beta-catenin upon the perturbation. Trajectories are displayed from the beginning of the perturbation until the end of the perturbation (1000 timesteps). (B and C) Trajectories of all variables that still evolve after the perturbation is released, example of repetition n°66 of the perturbation. After t= 1000 timesteps, the perturbation is released. In repetition n°66, the value (global activity) of 17 variables, among 62, still evolves for another several timesteps to adjust to the new state (without perturbation) before the system reaches a final stable state. Panel B displays the 9 first variables and panel C the 7 others. All other variables remained stable after releasing the perturbation. (D and E) time evolution of SOX9, Col-II and P38 (for panel D) and Beta-catenin, NFkB and RUNX2 (for panel E) under EPHA2 and inflammatory cytokine activation. Continuous lines denote the average global activity of each variable over the 100 repetitions, dashed lines denote + and – standard deviation over the 100 repetitions.

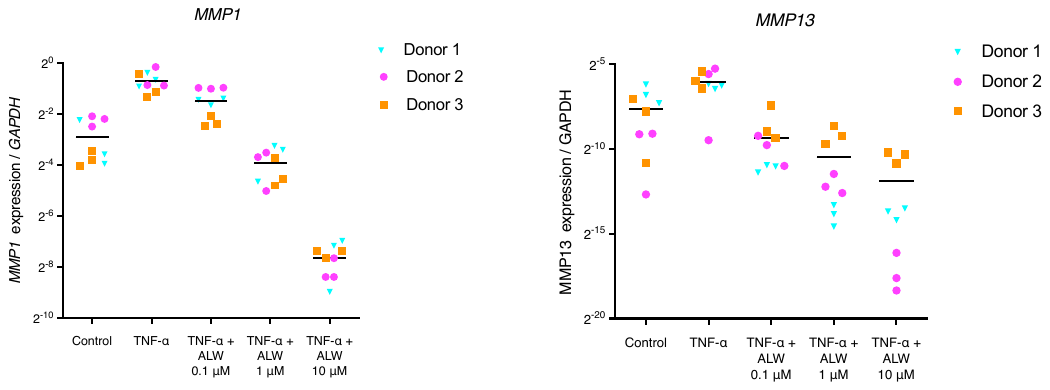

**Supplementary figure 3.** ALW-II-41-27 attenuates the expression of catabolic enzymes in OA cartilage explants in a dose-dependent manner.

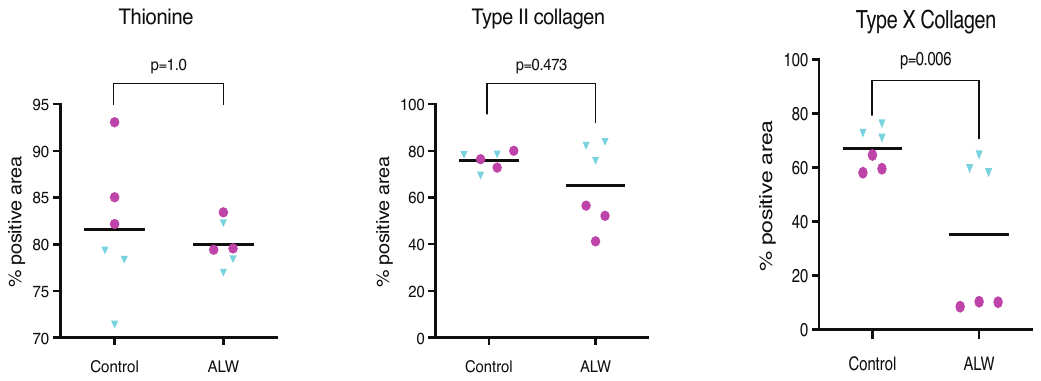

**Supplementary figure 4.** ALW-II-41-27 decreases type X collagen deposition but did not affected type II collagen and glycosaminoglycans. Percentage of positive area of thionine, type II collagen and type X collagen of tissue engineered cartilage derived from MSCs. Donors are represented with different colors and symbols: violet circles (donor 1) and blue triangles (donor 2). The horizontal line in the graphs represents the mean. For statistical analysis, the linear mixed model with Bonferroni’s multiple comparisons test was performed.

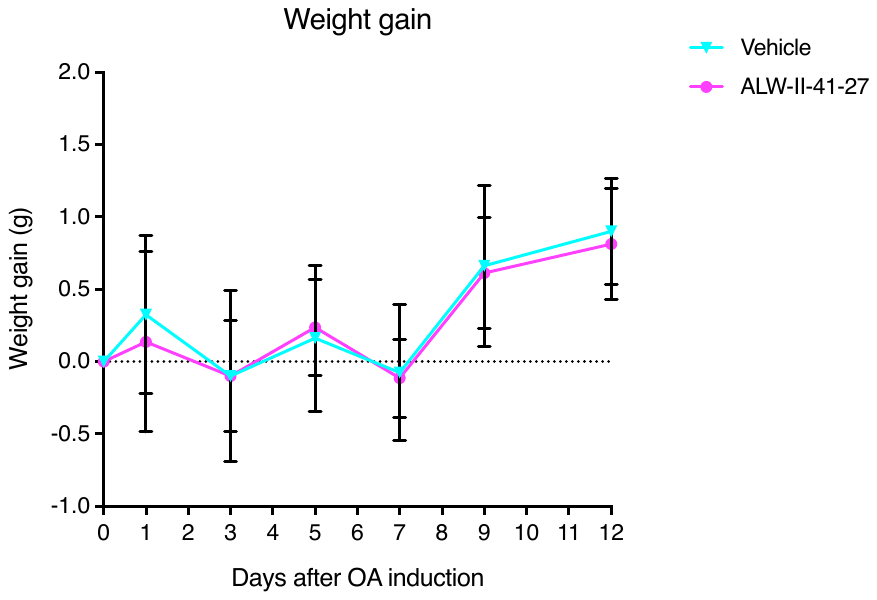

**Supplementary figure 5.** Weight gain of mice. Total body weight of each mice was determined on day 0, 1, 3, 5, 7, 9 and 12 using a scale. The average weight of the experimental group, vehicle- or ALW-II-41-27-treated mice (n=8) is shown.

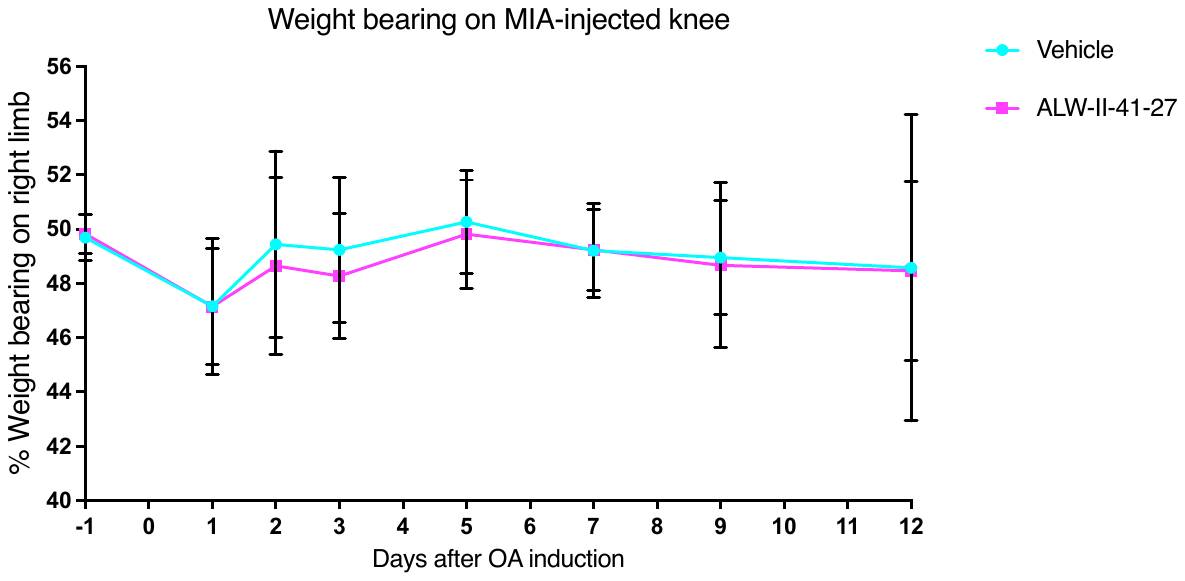

**Supplementary figure 6.** Pain measurements with incapacitance tester. Weight bearing between limbs along the study showed no differences between treated and vehicle group

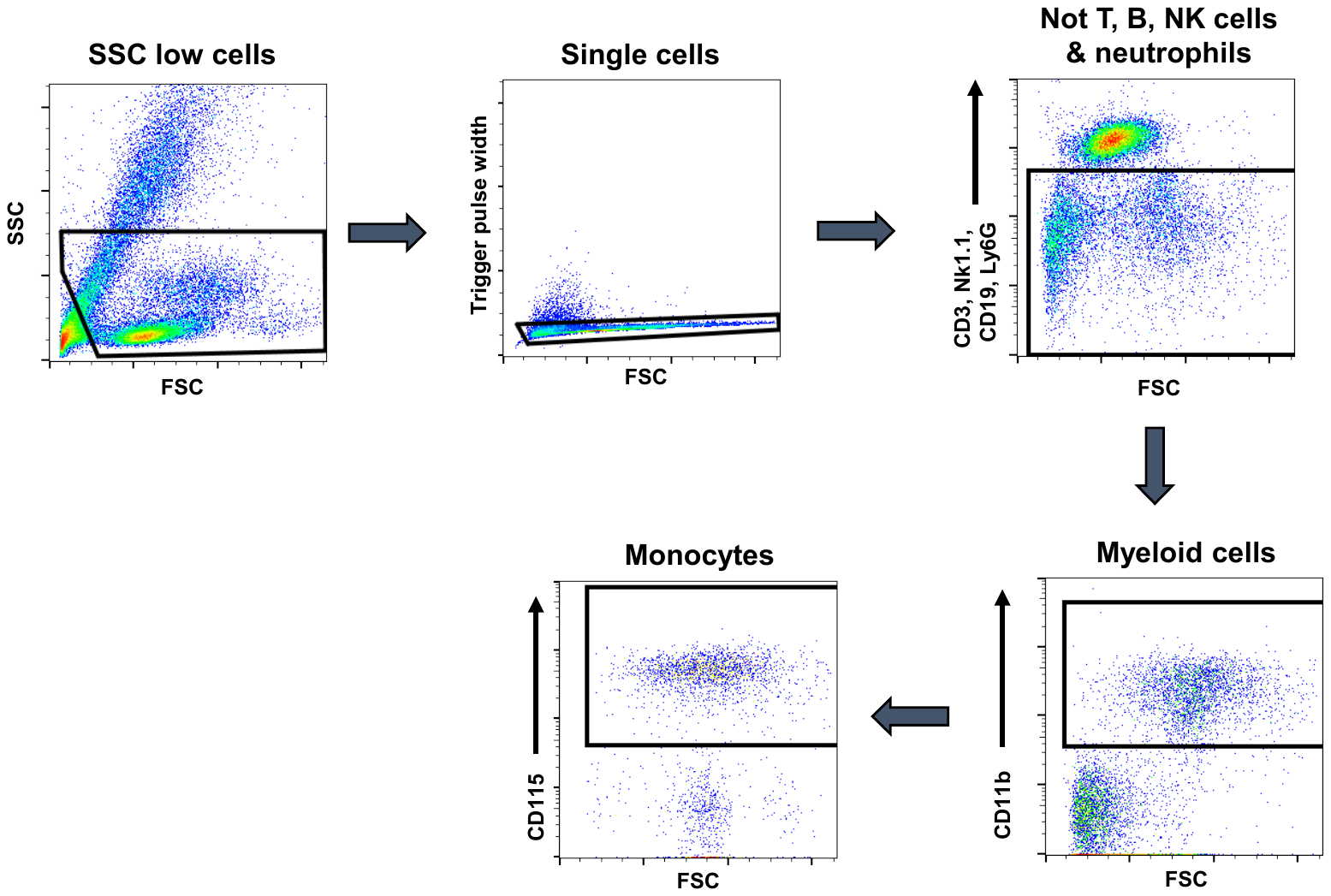

**Supplementary figure 7.** Gating strategy applied for blood monocytes analysis

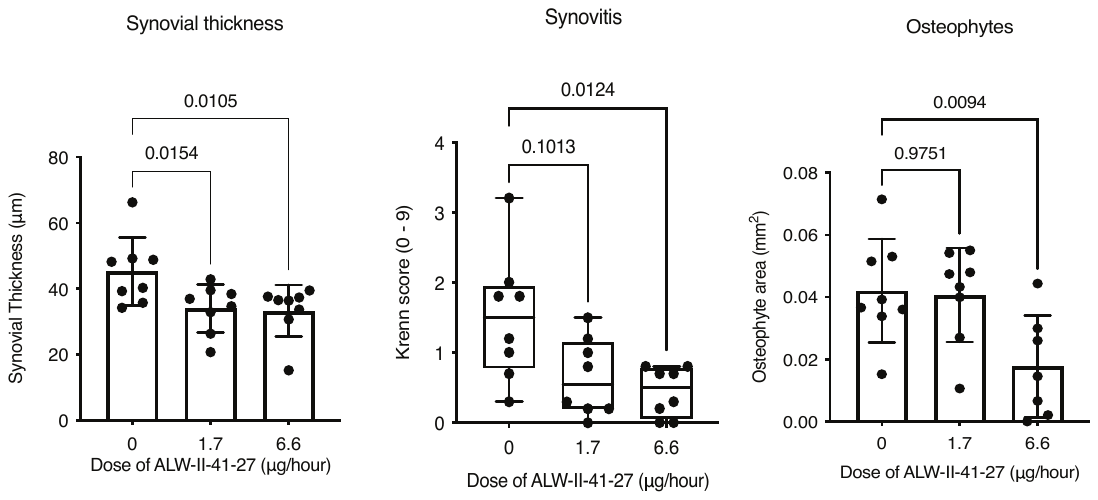

**Supplementary figure 8.**  ALW-II-41-27 treatment attenuates synovitis and osteophytosis. Analysis of synovial thickness, Krenn score and osteophyte area adjacent to the lateral side of the patella in mice containing an osmotic pump with vehicle (dose 0 μg/ hour) or ALW-II-41-27 (dose 1.7 and 6.6 μg/ hour). The data for the vehicle and the dose 6.6 μg/ hour is already shown in figure 5 and 6. Each dot represents data of an individual mouse (n=8). For statistical analysis, the linear mixed model with Bonferroni’s multiple comparisons test was performed
