## Supplementary Tables for "In silico, in vitro, and in vivo models reveal EPHA2 as a target for decreasing inflammation and pathological endochondral ossification in osteoarthritis"

| **Protein signaling** | | | |
| --- | --- | --- | --- |
| Source | Target | Sign | Reference |
| EPHA2 | Dsh | 1 | 2 |
| EPHA2 | ERK1/2 | 1 | 5 |
| EPHA2 | PI3K | 1 | 6 and 7 |
| AKT | EPHA2 | 1 | 4 |
| WNT | EPHA2 | 1 | 1 |
| **Genetic regulation** | | | |
| Source | Target | Sign | Reference |
| NFKB | EPHA2 | 1 | 3 |
| RAS | EPHA2 | 1 | 8 |

1	Peng Q, Chen L, Wu W, Wang J, Zheng X, Chen Z, et al. EPH receptor A2 governs a feedback loop that activates Wnt/beta-catenin signaling in gastric cancer. Cell Death Dis. 2018 Nov 19; 9(12):1146.

2	Li JY, Xiao T, Yi HM, Yi H, Feng J, Zhu JF, et al. S897 phosphorylation of EphA2 is indispensable for EphA2-dependent nasopharyngeal carcinoma cell invasion, metastasis and stem properties. Cancer Lett. 2019 Mar 1; 444:162-174.

3	Irie N, Takada Y, Watanabe Y, Matsuzaki Y, Naruse C, Asano M, et al. Bidirectional signaling through ephrinA2-EphA2 enhances osteoclastogenesis and suppresses osteoblastogenesis. J Biol Chem. 2009 May 22; 284(21):14637-14644.

4	Miao H, Li DQ, Mukherjee A, Guo H, Petty A, Cutter J, et al. EphA2 mediates ligand-dependent inhibition and ligand-independent promotion of cell migration and invasion via a reciprocal regulatory loop with Akt. Cancer Cell. 2009 Jul 7; 16(1):9-20.

5	Pratt RL, Kinch MS. Activation of the EphA2 tyrosine kinase stimulates the MAP/ERK kinase signaling cascade. Oncogene. 2002 Oct 31; 21(50):7690-7699.

6	Pandey A, Lazar DF, Saltiel AR, Dixit VM. Activation of the Eck receptor protein tyrosine kinase stimulates phosphatidylinositol 3-kinase activity. J Biol Chem. 1994 Dec 2; 269(48):30154-30157.

7.	Kim HS, Won YJ, Shim JH, Kim HJ, Kim BS, Hong HN. Role of EphA2-PI3K signaling in vasculogenic mimicry induced by cancer-associated fibroblasts in gastric cancer cells. Oncol Lett. 2019 Sep; 18(3):3031-3038.

8	Macrae M, Neve RM, Rodriguez-Viciana P, Haqq C, Yeh J, Chen C, et al. A conditional feedback loop regulates Ras activity through EphA2. Cancer Cell. 2005 Aug; 8(2):111-118.

**Supplementary table 1 : List of regulatory interactions between EPHA2 and other components of the network.** Each regulatory link is directed from the Source to the Target entity and the Sign indicates the type of influence (1 stands for activation). Literature references are provided to support the assumptions.

| **Variable index** | **Variable name** |
| --- | --- |
| 1 | WNT |
| 2 | DSH |
| 3 | IFG-I |
| 4 | R-SMAD |
| 5 | IHH |
| 6 | GLI2 |
| 7 | β-CATENIN |
| 8 | LEF/TCF |
| 9 | RUNX2 |
| 10 | SOX9 |
| 11 | PTHRP |
| 12 | PPR |
| 13 | COL-X |
| 14 | PKA |
| 15 | MEF2C |
| 16 | FGF |
| 17 | FGFR3 |
| 18 | STAT1 |
| 19 | SMAD complex |
| 20 | COL II |
| 21 | NKX3.2 |
| 22 | ERK1/2 |
| 23 | TGFβ |
| 24 | MMP13 |
| 25 | SMAD7 |
| 26 | SMAD3 |
| 27 | FGFR1 |
| 28 | ATF2 |
| 29 | NFκβ |
| 30 | HDAC4 |
| 31 | CCND1 |
| 32 | DLX5 |
| 33 | BMP |
| 34 | P38 |
| 35 | GSK3β |
| 36 | DC |
| 37 | PP2A |
| 38 | AKT |
| 39 | PI3K |
| 40 | ETS1 |
| 41 | RAS |
| 42 | IGF-IR |
| 43 | MSX2 |
| 44 | δEF-1 |
| 45 | ATF4 |
| 46 | HIF-2α |
| 47 | GREM1 |
| 48 | DKK1 |
| 49 | FRZB |
| 50 | Frizzled-LRP5/7 |
| 51 | Cytokines |
| 52 | ALK1 |
| 53 | ALK5 |
| 54 | Receptor inflam |
| 55 | TAK1 |
| 56 | JNK |
| 57 | Proteoglycans |
| 58 | IkB-alpha |
| 59 | SOCS |
| 60 | FOXO1 |
| 61 | IGFBP |
| 62 | EPHA2 |

**Supplementary table 2: List of in silico variables:** molecular entities included in the model and index of the corresponding variables as used in mathematical equations.

| Antibody | Clone | Fluorophore |
| --- | --- | --- |
| Anti-mouse/human CD11b | M1/70 | PerCp-Cy5.5 |
| Anti-mouse CD115 | AFS98 | PE |
| Anti-mouse Ly6C | HK1.4 | FITC |
| Anti-mouse CD62L | MEL-14 | APC |
| Anti-mouse Ly6G | 1A8 | PE-Cy7 |
| Anti-mouse CD3 | 17A2 | PE-Cy7 |
| Anti-mouse NK1.1 | PK136 | PE-Cy7 |
| Anti-mouse CD19 | 6D5 | PE-Cy7 |
| Anti-mouse F4/80 | BM8 | FITC |
| Anti-mouse CD86 | GL-1 | PE-Cy7 |
| Anti-mouse CD206 | C068C2 | APC |
| Anti-mouse CD163 | TNKUPJ | PE |
| Anti-mouse CD31 | 390 | APC |

**Supplementary Table 3**. Flow cytometry antibodies used for markers of interest
