## Supplementary Methods for "In silico, in vitro, and in vivo models reveal EPHA2 as a target for decreasing inflammation and pathological endochondral ossification in osteoarthritis"

**Data mining of microarray datasets**

Three previously published datasets were leveraged for the purpose of the current study. To identify a target associated with chondrocyte hypertrophy, we used a dataset previously generated in different zones of the growth plate of 14 days old female Swiss White mice [1], using a 44k whole genome oligo microarrays (G4122A; Agilent Technologies, Santa Clara, CA, United States). A list of differently expressed genes between the proliferative (PR) and hypertrophic (H) layer was obtained from the supplementary data of the publication, generated via paired t-test and selected factors with a cut off value of 3-fold change in the expression of genes. To identify a target associated with osteoarthritis, we used a dataset previously generated in 10-week-old male C57BL/6 mice by surgical destabilization of the medial meniscus (DMM) [2] using a 44k whole genome oligo microarrays (G4122A; Agilent Technologies). A list of differently expressed genes from cartilage of DMM- or sham-operated mouse joints was obtained from the supplementary data of the publication, generated via paired t-test and selected factors with a cut off value of 2-fold change in the expression of genes. To identify targets associated with human osteoarthritic cartilage we used a microarray dataset previously generated [3], performed using Affymetrix oligonucleotide microarray HG-U133plus2.0 (Affymetrix, Santa Clara, CA). Differently expressed genes in articular cartilage from OA and healthy donors were obtained from the supplementary data of the publication, generated via paired t-test and considering genes that displayed a mean fold change greater than 2. An online resource from Gene Ontology (GO) was used to obtain a list of genes associated with inflammation, namely GO: 0006954 Inflammatory response [4, 5]. The selected lists of uniquely expressed factors were then overlapped utilizing the Funrich software [6], to select for genes associated with osteoarthritis, inflammation and chondrocyte hypertrophy.

***In silico* experiments (simulations)**

A computational model of the intracellular signaling pathways regulating articular chondrocyte phenotypes [7, 8], was leveraged and completed with information about EPHA2. This mechanistic model represents 62 molecular entities (growth factors, receptors, transcription factors, extracellular matrix proteins and signaling) and the way they regulate each other, at protein and genetic level, to transduce external signals and define new cellular states (see list of variables in Supplementary Table S2 and the full list of interactions, in the standard .csv format in Supplementary Data 1 or at [insert GitHub link]). Connections of EPHA2 with the rest of the signaling network were added according to information found in literature (Fig 2A and supplementary table 1) with supporting references featuring different cell types, given that EPHA2 is still understudied in cartilage. The mathematical implementation consists of additive equations in which each variable (i.e. biomolecule) is regulated at protein (fast) and genetic (slow) level with values continuously ranging between 0 (fully repressed) and 1 (fully expressed and activated). The global activity is defined as the multiplication of the genetic activation level and protein activation level. An asynchronous updating scheme with priority classes (i.e. fast sub-variables updated first) was employed, resulting in semi-quantitative stochastic simulations [9]. In the algorithm executing the asynchronous updating scheme to simulate the model, the fast (protein signaling level) and slow (genetic level) sub-variables are updated one-by-one with two different priority classes. A new slow sub-variable is updated when all fast sub-variables have been updated and when a pseudo-stable state has been reached at the fast (protein signaling) level. The order in which fast and slow sub-variables are updated within a priority class is random (MATLAB random seed via randperm() function). The algorithm counts a new computational timestep each time a new slow sub-variable is updated, this is how time-trajectories are obtained. The outcome of a given perturbation may vary, due to the random order in which variables are updated. This reflects actual biochemical processes, in which there is a random component for two molecules to enter in contact and react together/influence each other in the cytoplasm or nucleus, due to inherent entropy. Hence, each perturbation has a certain propensity to trigger a state transition. Therefore, each perturbation is repeated 100 times in order to compute the percentage of repetition triggering a state transition. The only parameter of the model is a saturation constant affecting the weight of a regulatory interaction in the network. The saturation constant was assigned an arbitrary value (s=2/3) based on a previous study which evaluated the influence of that constant on a similar model [9]. This constant determines how fast a protein activity or gene expression can saturate to the maximal value depending on the amount of excess positive and negative upstream interactions.

A Monte Carlo approach (10,000 random initializations) was used to compute the model’s stables states, equating to possible molecular states of the cell (i.e. chondrocyte phenotypes). The stable states were characterized regarding the variable (i.e. biomolecules’ activity) as markers. In particular, the states with a high (resp. low) activity for SOX9 and low (resp. high) activity for RUNX2 where considered as healthy (resp. hypertrophic) chondrocytes. *In silico* experiments were initially executed using the model’s stable state resembling the most a regular healthy chondrocyte. *In silico* experimental conditions were applied by (simultaneously) forcing the targeted variable(s) to be set to 0 (for inhibition) or 1 (for activation), unless specified otherwise (e.g. variation of EPHA2 activity in Fig. 2B). The *in silico* conditions were applied for 1,000 computing steps after which all variables were left free to evolve until reaching a new stable state, thereby simulating a bolus treatment effect. This was repeated 100 times and the various outcomes, or final states, (due to the model stochasticity) were averaged to compute the final profiles, standard deviations were also computed. When a final state is different from the initial state before perturbation, this is called a state transition. The percentage of transition to each emerging final state was also computed over the 100 repetitions. The period during which the perturbation was maintained was set to 1000 timesteps, as it is largely greater than the time typically needed (about 400steps) for all variables to reach a steady state in the perturbed scenarios. The following 4 *in silico* conditions (or perturbations) were applied: (1) activation of EPHA2 (variable #62), (2) activation of pro-inflammatory cytokines(variable #51), (3) the combination of the two previous conditions and (4) blockage of EPHA2 while activating the pro-inflammatory cytokines. The model and associated code are available via the following GitHub repository: https://github.com/Rapha-L/Virtual_Chondrocyte_for_EPHA2_study

**Evaluation of ALW-II-41-27 in OA chondrocytes and cartilage explants**

Human articular cartilage was obtained with implicit consent as waste material from patients undergoing total knee replacement surgery. This protocol was approved by the Medical Ethical Committee of the Erasmus MC, University Medical Center, Rotterdam, protocol number MEC-2004-322. Full thickness cartilage explants (ø = 5 mm) were harvested from macroscopically intact areas and washed twice with 0.9% NaCl (Sigma Aldrich, St. Louis, MO, USA). To isolate chondrocytes, cartilage chips were subjected to protease (2 mg/mL, Sigma Aldrich) for 2 hours followed by overnight digestion with 1.5 mg/mL collagenase B (Roche Diagnostics, Basel, Switzerland) in Dulbecco’s modified Eagle’s medium (DMEM) high glucose supplemented with 10% fetal bovine serum. Single cell suspension was obtained by filtrating the cellular solution by a 100 µm filter. The isolated chondrocytes were expanded in monolayer at a seeding density of 7,500 cells/cm2 in DMEM high glucose supplemented with 10% fetal bovine serum, 50 μg/mL gentamicin, and 1.5 μg/mL fungizone (Gibco, Grand Island, NY, USA). Approximately 80% confluency cells were trypsinized and reseeded at 7,500 cells/cm2. Cells were used for experiments after 3 passages. To re-differentiate the expanded articular chondrocytes, a 3D alginate bead culture model was used [10, 11]. Alginate beads were prepared by mixing passage three chondrocytes and re-suspended them in 1.2% (w/v) low viscosity alginate (Kelton LV alginate, Kelko Co, San Diego, CA, USA) in 0.9% NaCl (Sigma Aldrich) at a concentration of 4 × 10^6^ cells/mL. Beads were made by dripping the cell-alginate suspension in 105 mM CaCl_2_ (Sigma Aldrich) through a 22-gauge needle. Beads were washed with 0.9% NaCl and DMEM low glucose. Beads with a size that deviated from the average after a visual inspection were not included in the experiment. Re-differentiation of chondrocytes was performed in a 24-well plate (BD Falcon) for two weeks in 100 μL/bead DMEM low glucose supplemented with 1% v/v Insulin-Transferrin-Selenium (ITS™+ Premix, Corning, Bedford, MA, USA), 10 ng/ml transforming growth factor beta-1 (TGF-β-1, recombinant human, R&D systems) 25 μg/mL l-ascorbic acid 2-phosphate (Sigma Aldrich), 50 μg/ml gentamicin, and 1.5 μg/mL fungizone (both Gibco). After two weeks, TGF-β-1 was no longer added to the medium and cells were cultured with 10 µM of ALW-II-41-27 (MedChemExpress, Ann Arbor, MI, USA), vehicle (DMSO) and/or with the pro-inflammatory cytokine TNF-α 10 ng/mL for 24 hours. Preliminary experiments were performed using two doses of ALW-II-41-27, 1 and 10 µM, being 10 µM more effective. Medium and alginate beads were harvested for further analyses.

**Evaluation of ALW-II-41-27 in chondrogenically differentiated MSCs**

Human MSCs were isolated from surplus iliac crest bone chip material harvested from pediatric patients undergoing alveolar bone graft surgery. All human samples were obtained with the approval of the Erasmus MC, University Medical Center Medical Research Ethics Committee (MEC-2014-16). Written consent was not required in accordance with the national code regarding the use of waste surgical material for scientific research (www.coreon.org), and an opt-out option was available. Iliac crest bone chips were washed with expansion medium composed of Minimum Essential Medium (MEM)-α (containing nucleosides) supplemented with heat inactivated 10% v/v fetal bovine serum (FBS) (both Thermo Fisher Scientific, Waltham, MA, USA), 1.5 µg/ml fungizone (Gibco), 50 µg/ml gentamicin (Gibco), 25 µg/ml L-ascorbic acid 2-phosphate (Sigma-Aldrich, St. Louis, MO, USA), and 1 ng/ml fibroblast growth factor-2 (Instruchemie, Delfzijl, The Netherlands), and the resulting cell suspension was seeded in T75 flasks. Cells were washed twice with phosphate buffered saline (Thermo Fisher Scientific) supplemented with 2% v/v heat inactivated FBS 24 h following seeding to remove non-adherent cells. MSCs were cultured at 37°C and 5% carbon dioxide under humidified conditions, with expansion medium refreshed every 3–4 days. MSCs were sub-cultured upon reaching 80–90% confluency using 0.25% w/v trypsin-EDTA (Thermo Fisher Scientific) and reseeded at a cell density of 2,300 cells/cm^2^. MSCs were used at passage 3 for chondrogenic pellet cultures.

For chondrogenic differentiation, 2 × 10^5^ MSCs were suspended in 500 µl of chondrogenic differentiation medium composed of high glucose Dulbecco’s Modified Eagle Medium supplemented with 1.5 µg/ml fungizone (Gibco), 50 µg/ml gentamicin (Gibco), 1 mM sodium pyruvate (Thermo Fisher Scientific), 1% v/v Insulin-Transferrin-Selenium (ITS™+ Premix, Corning, Bedford, MA, USA), 40 µg/ml proline (Sigma-Aldrich), 25 µg/ml L-ascorbic acid 2-phosphate (Sigma-Aldrich), 100 nM dexamethasone (Sigma-Aldrich), and 10 ng/ml TGF-β-1 (R&D systems). The cell suspension was added to 15 ml conical polypropylene tubes (TPP, Radnor, PA, USA) and centrifuged at 300 g for 8 min to facilitate pellet formation. Chondrogenic MSC pellets were cultured at 37°C and 5% carbon dioxide in a humidified atmosphere. After 24 h, pellets were tapped to enhance pellet formation and the medium was renewed with chondrogenic medium with 100 nM of ALW-II-41-27 (MedChemExpress, Ann Arbor, MI, USA) or vehicle (DMSO). Afterwards the medium was renewed two times per week for a period of 3 weeks.

**Nitric oxide (NO) assay**

NO production was measured in the medium of OA chondrocytes by determining the content of nitrite using the Griess reagent (Sigma Aldrich). The reaction was monitored at 540 nm using a spectrophotometer (VersaMax; Molecular Devices, Sunnyvale, USA). Sodium nitrite (NaNO_2_; Chemlab, Zedelgem, Belgium) was used as standard for the calibration curve.

**Interleukin-6 assay**

A commercially available enzyme-linked immunosorbent assay (ELISA) kit was used to determine the concentration of IL-6 in the medium of OA chondrocytes as per manufacturer's instructions (R&D systems, Minneapolis, MN, USA).

**Histological analysis of chondrogenic pellets**

After 3 weeks of chondrogenic induction, pellets were fixed with 4% (v/v) formaldehyde in phosphate buffered saline, embedded in paraffin and sectioned (6 μm). Glycosaminoglycan (GAG) was stained with 0.04% thionine solution and collagen type II was immunostained using a primary antibody II-II6B3 (Developmental Studies Hybridoma Bank, Iowa City, IA, USA) 0.4 µg/mL in PBS/1% bovine serum albumin (BSA; Sigma-Aldrich), collagen type X using primary antibody 14-9771-82, 5 µg/mL in PBS/1% BSA (Thermofisher, Waltham, MA, USA). Antigen retrieval for collagen type II was performed with 1 mg/mL pronase (Sigma-Aldrich) in PBS for 30 min at 37°C, followed by incubation with 1% hyaluronidase (Sigma-Aldrich) in PBS for 30 min at 37°C to improve antibody penetration. Antigen retrieval for collagen type X was done by a 2h incubation in 1 mg/mL pepsin in 0.5M acetic acid followed by incubation with 1% hyaluronidase (Sigma-Aldrich) in PBS for 30 min at 37°C. The slides were pre-incubated with 10% normal goat serum (Sigma-Aldrich) in PBS with 1% BSA (Sigma-Aldrich). Next, the slides were incubated for 1 h (for collagen type II) or overnight (for collagen type X) with the primary antibody, and then with a biotin-conjugated secondary antibody (HK-325-UM, Biogenex, Fremont, CA, USA), alkaline phosphatase-conjugated streptavidin (HK-321-UK, Biogenex), and the Neu Fuchsin chromogen (B467, Chroma Gesellschaft). An IgG1 isotype antibody (X0931, Dako Cytomation, Santa Clara, CA, USA) was used as negative control.

**Gene expression**

Alginate beads were dissolved using citrate buffer, centrifuged at 200 g and the pellet was resuspended in RLT (Qiagen, Hilden, Germany) buffer containing 1% beta-mercaptoethanol for RNA isolation. RNA was isolated from the cartilage explants by snap freezing in liquid nitrogen followed by pulverization using a Mikro-Dismembrator (B. Braun Biotech International GmbH, Melsungen, Germany) at 2800 rpm. The tissue was homogenized with 18 μL/mg sample RNA-Bee TM (Tel-Test Inc., Friendswood, TX, USA) and 20% chloroform.

The MSCs chondrogenic pellets were homogenized in RNA-Bee TM (Tel-Test Inc., Friendswood, USA) and 20% chloroform was added to extract RNA. mRNA isolation was performed according to manufacturer’s protocol utilizing the RNeasy Column system (Qiagen, Hilden, Germany). The RNA concentration was determined using a NanoDrop spectrophotometer (Isogen Life Science, Utrecht, The Netherlands). 0.5 μg RNA was used for cDNA synthesis following the protocol of the manufacturer of the RevertAid First Strand cDNA kit (Thermo Fisher Scientific, Waltham, MA, United States). qPCR was performed on a Bio-Rad CFX96 Real-Time PCR Detection System (Bio-Rad) to assess gene expression, Alkaline phosphatase (*ALPL*, Fw: GACCCTTGACCCCCACAAT; Rev: GCTCGTACTGCATGTCCCCT; Probe: TGGACTACCTATTGGGTCTCTTCGAGCCA), Collagen type 2 (*COL2A1*; Fw: GGCAATAGCAGGTTCACGTACA ; Rev: CGATAACAGTCTTGCCCCACTT; Probe: CCGGTATGTTTCGTGCAGCCATCCT), Collagen type 10 (*COL10A1*; Fw: CAAGGCACCATCTCCAGGAA; Rev: AAAGGGTATTTGTGGCAGCATATT; Probe: TCCAGCACGCAGAATCCATCTGA), matrix metalloproteinase-13 (*MMP13*; Fw: AAGGAGCATGGCGACTTCT; Rev: TGGCCCAGGAGGAAAAGC; Probe: CCCTCTGGCCTGCGGCTCA), Runt-related transcription factor 2 (*RUNX2*; Fw: ACGTCCCCGTCCATCCA; Rev: TGGCAGTGTCATCATCTGAAATG; Probe: ACTGGGCTTCTTGCCATCACCGA), Tumor Necrosis Factor-a (*TNFA*; Fw: GCCGCATCGCCGTCTCCTAC; Rev: AGCGCTGAGTCGGTCACCCT). Glyceraldehyde-3-phosphate dehydrogenase (*GAPDH*; Fw: ATGGGGAAGGTGAAGGTCG; Rev: TAAAAGCAGCCCTGGTGACC; Probe: CGCCCAATACGACCAAATCCGTTGAC) was found stable and therefore used as reference gene. Data were analyzed by the ΔΔCt method and normalized to the expression of *GAPDH* of each condition and compared to the corresponding gene expression in the control groups.

**Western blot analysis**

Mesenchymal stromal cells were expanded to approximately 50 % confluence. Next, medium was refresh with α-MEM supplemented with 1 mg/ml bovine serum albumin (Sigma), 1.5 μg/mL Amphotericin B (Invitrogen), 25 μg/mL L-ascorbic acid 2-phosphate (Sigma-Aldrich), 50 μg/mL gentamycin (Invitrogen). The following day, cells were first exposed to 0,1 – 10 μM ALW-II-41-27 (Cayman Chemicals) for 1 hour and then treated with 10 ng/ml TNF-α and 0,1 – 10 μM ALW-II-41-27. After 30 minutes cells were washed with PBS on ice and incubated with mammalian cell lysis buffer, supplemented with 1% Halt Protease Inhibitor (Thermo Scientific) and 1% Halt Phosphatase Inhibitor (Thermo Scientific). Cells were manually scraped off the surface and cell lysate was harvested and stored at -20 °C. Samples were sonicated using a BioRuptor® Pico sonication device for 10 cycles of 30 seconds on and 30 seconds off. Samples were centrifuged to remove remaining cell debris; the protein concentration was determined using a BCA assay and protein contents were analysed by western blot. Proteins were denatured in reducing conditions and loaded onto a 4-12% sodium dodecyl sulfate-polyacrylamide gel (Thermo Fisher). 5-20 µg of protein was run through the gel at 120V for 1 hour. The gel was blotted onto a PVDF membrane at 20V for 1 hour using a ThermoFisher wet transfer system. The membranes were blocked using 5% milk in Tris buffer saline Tween (TBST) buffer for 1-3 hours. The membranes were washed with TBST and incubated with 5% BSA in TBST containing the primary antibodies for the proteins of interest overnight in the fridge. Next, the membranes were washed with TBST and incubated with the secondary antibody for 2-3 hours at room temperature. Afterwards, the membranes were washed with TBST and signal was visualized using a SuperSignal™ West Pico Detection Kit for rabbit IgG that produces a chemiluminescent signal when exposed to UV light. Pictures were developed using a Uvitec Alliance developer. Antibodies used: EphA2 (1:1000, Cell signalling, D4A2, Rabbit), phosphorylated-EphA2 (1:1000, Cell signalling, S897, Rabbit), alfa-Tubulin (1:1000, Cell signalling, 11H10, Rabbit).

**Animal model**

All animal experimentation procedures were conducted with approval by the Animal Ethical Committee of Erasmus University Medical Center (License number AVD101002015114, protocol number 16-691-06). 12-week-old male C57BL/6 mice (C57BL/6J0laHsd, 27.01 g ± 2.05 g; Envigo, Cambridgeshire, UK), were housed in groups of 8 in individually ventilated cages and maintained on a 12 h light/dark cycle with ad libitum access to standard diet and water at the Experimental Animal Facility of the Erasmus MC. Mice were randomly divided into two experimental groups (N=8 per group): Control and ALW-II-41-27-treated mice. For all procedures, mice were anesthetized using 3% isoflurane/0.8 L O_2_/min (Pharmachemie BV, Haarlem, the Netherlands). OA was induced unilaterally by intra-articular injections of 60 μg Monoiodoacetate (MIA) (Sigma-Aldrich, St. Louis, USA) in 6 μl of saline (0.9% NaCl; Sigma-Aldrich) at day 0. Injections were performed after a 3-4 mm dermal incision was made to the right knee at the height of the patellar tendon. All intra-articular injections were administered using a 50 µl syringe (Hamilton, Bonaduz, Switzerland) and 30G needle (BD Medical, New Jersey, USA). ALW-II-41-27 was delivered using Alzet micro-osmotic pumps (Durect Corporation, CA, USA) model 1004, delivery rates of 0.11 μL/ hour, that were implanted subcutaneously on the back of the mice, slightly posterior to the scapulae, immediately after the intra-articular injections. Osmotic pumps were filled with dimethyl sulfoxide: polyethylenglicol alone (55:45 ratio, vehicle-treated group, N=8 mice) or containing 6 mg of ALW-II-41-27 dissolved in vehicle (treated group, N=8 mice) which leads to a dose of 6.6 μg/ hour. A third group of N=8 mice received the implantation of osmotic pumps delivering a dose of 1.7 μg/ hour of ALW-II-41-27. In the figures 5 and 6 we report the 6.6 μg/ hour dose of ALW-II-41-27. Synovial thickness, Krenn score and osteophyte size for all groups can be found in supplementary figure 9. Mice were euthanized in agreement with the Directive 2010/63/EU by cervical dislocation under isoflurane anesthesia 14 days following MIA injection. After which knees were fixed in 4 % formalin (v/v) for 1 week, decalcified in 10 % EDTA for 2 weeks and embedded in paraffin. Coronal sections of 6 μm were cut for analyses.

**Flow cytometric analysis of peripheral blood monocytes**

Peripheral blood was harvested from the facial vein of mice on days 2- and 12 post-induction of OA, as previously described [12]. Blood sampling order was performed randomly at each time-point. 50 µl of whole blood was pre-incubated with purified rat anti-mouse CD16/CD32 (BD Biosciences Cat# 553140, New Jersey, USA) for 5 mins on ice. Blood was stained for the expression of CD11b (BioLegend, San Diego, USA, Cat# 101228), CD115 (BioLegend, Cat# 135505), Ly-6C (BioLegend Cat# 128005) and CD62L (BioLegend, Cat# 104412) to identify myeloid cells and specific monocyte subsets, as well as CD3 (BioLegend, Cat# 100220), NK1.1 (BioLegend, Cat# 108713), CD19 (BioLegend, Cat# 115520) and Ly-6G (BioLegend, Cat# 127618) to eliminate T cells, natural killer cells, B cells and neutrophils (Supplementary table 3). Cells were stained for 30 mins at 4°C in the dark, followed by incubation with 2 ml of 1X FACS lysing solution (BD Biosciences) for 10 minutes to lyse red blood cells. Following centrifugation at 400 g for 10 minutes, supernatant was removed and cells washed and resuspended in FACSFlow buffer (BD Biosciences).

All samples were analyzed using a FACSJazz cytometer (BD Biosciences) and FlowJo software version 10.0.7 (FlowJo LLC, Oregon, USA). The gating strategies applied for blood monocytes analysis are presented in supplementary figure 8.

**Histological analyses of murine knee joints**

To evaluate cartilage damage, sections were stained with Safranin O & Fast Green. To evaluate synovial inflammation and osteophyte area, sections were stained with Hematoxylin & Eosin. Images were acquired using the NanoZoomer Digital Pathology program (Hamamatsu Photonics, Ammersee, Germany). For each knee, 3 sections of the patellofemoral compartment, taken at standardized locations in the knee with 180 μm distance in between were evaluated by two independent well trained evaluators that were blinded for the treatment (MNFB and NK). For each knee, the average value of the three sections was calculated and the values of two observers were averaged and used for representation and statistical analyses.

Cartilage damage was evaluated with the Osteoarthritis Research Society International (OARSI) scoring system [13].

Osteophyte size was assessed by measuring the area of the osteophyte at the lateral side of the patella, the location where the incidence of osteophytes was highest, using the NanoZoomer digital pathology program.

Synovial thickness was measured from the capsule to the superficial layer of the synovial membrane at the medial and lateral sides of the parapatellar recesses (three positions per section).

Synovitis was evaluated using the Krenn score [14] which considers three features of chronic synovitis (enlargement of lining cell layer, cellular density of synovial stroma, leukocytic infiltrate) and grade each feature from 0 (absent) to 3 (strong). The sum provided the synovitis score, which is interpreted as follows: 0–1, no synovitis; 2–4, low-grade synovitis; 5–9, high-grade synovitis.

To evaluate macrophages in the synovial membrane, F4/80 was used as a marker. For this purpose, an antigen retrieval was performed by placing the slides in a 20 µg/mL proteinase K solution for 30 minutes at 37°C. Blocking of specific binding was performed with 10 % goat serum (Southern Biotech, Birmingham, AL, USA) for 30 min. Hereafter, sections were incubated for 1 h with 1 µg/mL primary antibody F4/80 (eBioscience #14-4801-82, Waltham, MA, USA), followed by 30 min incubation with a biotinylated rabbit anti-rat IgG l (Vector, BA-4000, Burlingame, CA, USA) 6 µg/mL in PBS/1 % BSA. Thereafter, sections were incubated with an alkaline-phosphatase- conjugated streptavidin label (HK-321-UK, Biogenex) concentration not stated, diluted 1:50 in PBS/1 % BSA. To reduce background, endogenous alkaline phosphatase activity was inhibited with levamisole (Sigma-Aldrich Chemie). New Fuchsin (Fisher Scientific) and Napthol AS-MX phosphate (Sigma-Aldrich Chemie) substrates were used for color development and counterstaining was performed with hematoxylin. As a negative control, a rat IgG2a antibody (#14-4321-82, eBioscience Inc. San Diego, CA, USA) was used.

To evaluate type X collagen in mice knees, a pre-coupling step was performed 24 hours before the staining using a 1:10 dilution of collagen type X antibody (Quartett, #X53) with fluorescent goat anti-rat IgG (H+L) (Cross-Adsorbed Secondary Antibody, Alexa Fluor 546, #A11081, Fisher Scientific, Landsmeer, The Netherlands). Mouse IgG1 antibody (Dako Cytomation #X0931) dilution 1:10, was used as a negative control. Antigen retrieval was performed using Pepsin (Sigma #P7000) 1 mg/mL in 0.5M acetic acid pH2 for 2 hours following 10 mg/ml hyaluronidase (Sigma #H3506) for 30 minutes. Slides were incubated with 10% normal goat serum (Southern Biotech #0060-01) for 30 minutes. Slides were incubated overnight with the pre-coupled antibodies and following day slides were mounted using ProLong Diamond Antifade Mountant with DAPI (#P36966, Fisher Scientific, Landsmeer, The Netherlands).

**Pain measurement: hind limb weight distribution**

Hind limb weight distribution was monitored as a surrogate pain indicator using an Incapacitance Tester (Linton Instrumentation, Norfolk, UK). Mice were positioned on the Incapacitance Tester with each hind limb resting on a separate force plate. Animals were habituated to the apparatus, three times per week, starting 2 weeks prior to the experiments. The examiner performing the measurements was blinded for treatment condition (MNFB). A baseline measurement was performed at day – 1, just before the induction of the OA. Follow up measurements were performed at day 1, 2, 3, 5, 7, 9 and 12. For data analyses, measurements with a registration below 10 g (< 30 % of total body weight) in total on both hind limbs were excluded. 10 measurements were recorded per mouse per time point, of which at least 7 measurements were available on average. For each time point per mouse, the average of these measurements was used to calculate the percentage of weight on the affected limb as an indication of pain in the affected limb.

**Data and statistical analyses**

Statistical evaluation was performed using GraphPad Prism 9.0 and IBM SPSS 24 (IBM). Each *in vitro* experiment included at least 3 biological replicates and was repeated with cells derived from 3 donors. The *in vivo* study was designed to generate groups of equal size, used randomization and blinded analyses. The declared group size is the number of independent values that were used for statistical analysis. Sample size for the *in vivo* study was calculated considering the weight distribution over the hind limbs, as readout parameter. Based on previous studies, we consider an increase of 13% (standard deviation of 10%) in weight distribution on the affected limb in time in the therapy groups as relevant in our study [15]. Sample size was calculated with a statistical power of 80% and significance level of 0.05, which led to N=8. For statistical analysis, the linear mixed model with Bonferroni’s multiple comparisons test was performed.
